# ACAD10 encodes two orphan enzymes in the ether lipid biosynthetic and salvage pathways

**DOI:** 10.1101/2025.09.10.675409

**Authors:** James S. Ye, Elena Purlyte, Victor A. Lopez, Jericha Mill, Lexus Tatge, Thomas Kizzar, Dominique Baldwin, Edrees Rashan, Juhee Kim, David J. Pagliarini, Diana R. Tomchick, Krzysztof Pawłowski, Peter M. Douglas, Judith Simcox, Vincent S. Tagliabracci

## Abstract

Ether lipids play critical roles in membrane dynamics, antioxidant defense, and signaling. They comprise ∼20% of mammalian phospholipids, and disruptions in their metabolism cause severe genetic disorders and are associated with neurodegenerative and metabolic diseases. Ether lipids are synthesized *de novo* from glycolytic intermediates or salvaged from the diet. While the products of these pathways are known, several key enzymes remain unidentified, including the 1-O-alkylglycerol kinase and the 1-O-alkyl-2-acetyl-*sn*-glycero-3-phosphate phosphatase. Here, we show that acyl-CoA dehydrogenase member 10 (ACAD10) catalyzes the phosphorylation of 1-O-alkylglycerols and the dephosphorylation of 1-O-alkyl-2-acetyl-*sn*-glycero-3-phosphate. Worms and mice lacking ACAD10 have reduced ether lipid levels and cannot salvage dietary alkylglycerols. Furthermore, individuals from the Akimel O’odham (Pima) tribe carrying *ACAD10* polymorphisms also show decreased plasma ether lipid levels. Collectively, our findings resolve two long-standing gaps in ether lipid biochemistry and reveal a mechanistic link between ether lipid metabolism and a population-associated risk factor for type 2 diabetes.

## Introduction

Ether lipids are a distinct subclass of glycerophospholipids, characterized by the substitution of the fatty acyl chain at the *sn*-1 position of the glycerol backbone with a fatty alkyl chain. Linked by an ether bond rather than the typical ester bond (**Figure 1A**), they are further classified as alkyl-ether or alkenyl-ether lipids, depending on whether the bond is an ether or a vinyl-ether^1,2^. Platelet-activating factor, for example, has an alkyl group at the *sn*-1 position and an acetyl group at the *sn*-2 position of the glycerol backbone^3^. In contrast, plasmalogens, the most prevalent ether lipids, have an alkenyl-ether bond at the *sn*-1 position and a long-chain acyl group at the *sn*-2 position^4^.

**Figure 1.**
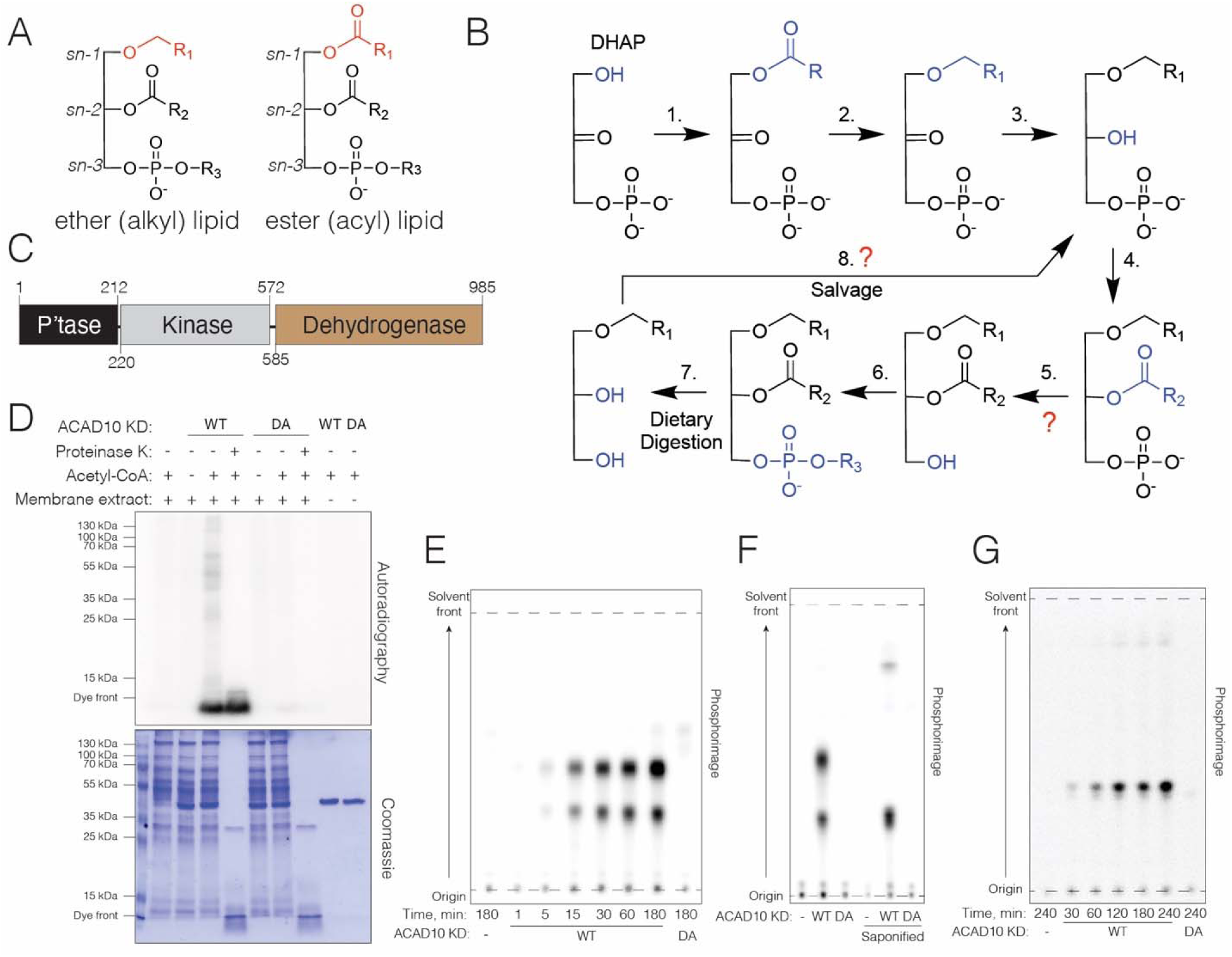
*C. elegans* ACAD10 phosphorylates 1-O-alkylglycerols *in vitro*. **(A)** Chemical structures of ether- and ester-linked glycerophospholipids. **(B)** Ether lipids are synthesized from dihydroxyacetone phosphate (DHAP), an intermediate of glycolysis. In step (**1**), DHAP is acylated by GNPAT, and then AGPS replaces the acyl group with a fatty alcohol, forming the ether linkage (**2**). Step (**3**) involves the reduction of the *sn*-2 ketone by AYR/PexRAP, followed by long or short chain acylation at *sn*-2 by AGPAT enzymes (**4**). During PAF biosynthesis, an unidentified alkylacetylglycerophosphatase catalyzes dephosphorylation of 1-O-alkyl-2-acetyl-*sn*-glycero-3-phosphate. During plasmalogen biosynthesis, phosphatidic acid phosphatases, such as LIPIN, catalyzes the dephosphorylation (**5**). Subsequently, choline/ethanolamine phospho-transferases add the headgroups in step (**6**). After dietary ether lipids are enzymatically digested (**7**), the acyl and phosphate head groups are removed, leaving the ether linkage intact. The resulting 1-O-alkylglycerols can re-enter the ether lipid biosynthetic pathway through phosphorylation by an unknown kinase **(8).** Adapted from^19^. **(C)** Schematic representation of the *C. elegans* ACAD10 protein depicting the phosphatase (P’tase), kinase and dehydrogenase domains. **(D)** Incorporation of ^32^P from [γ-^32^P]ATP, into a Proteinase K resistant species in a membrane extract by *C. elegans* ACAD10 kinase domain (ACAD10 KD; residues 209-568) or the mutant D424A (DA). Reactions were performed in the presence of Mg^2+^, acetyl CoA and a mouse liver mitochondrial extract, and the products were resolved by SDS–PAGE and visualized by autoradiography (top) and Coomassie staining (bottom). Data are representative of at least three independent experiments. **(E)** Thin layer chromatogram depicting the time-dependent incorporation of ^32^P from [γ-^32^P]ATP into two lipid species by *C. elegans* ACAD10 KD. Reactions were performed in the presence of Mg^2+^, acetyl CoA and a mouse liver mitochondrial extract. Lipids from **(D)** were extracted, separated by TLC and visualized by phosphorimaging. Data are representative of at least three independent experiments. **(F)** Thin layer chromatogram depicting the lipid species phosphorylated by *C. elegans* ACAD10 KD in a mitochondrial extract following saponification. Products were analyzed as in **(E)**. Data are representative of at least three independent experiments. **(G)** Thin layer chromatogram depicting the incorporation of ^32^P from [γ-^32^P]ATP, into chimyl alcohol by *C. elegans* ACAD10 KD. Reaction products were separated by TLC and visualized by phosphorimaging. Data are representative of at least three independent experiments.

Ether lipids play crucial roles in various biological processes, including membrane dynamics, signal transduction, cellular responses to oxidative stress and ferroptosis^5,6^. They are widespread across most domains of life, and in mammals, they make up around 20% of the total phospholipid pool^4,7–10^. In humans, disruptions in ether lipid metabolism have been linked to a range of diseases, including coronary artery disease, rhizomelic chondrodysplasia punctata, Alzheimer’s disease, type 2 diabetes, and others^11–14^. Interestingly, supplementation with ether lipids and their precursors has shown promise as a therapeutic strategy to improve outcomes in these conditions^15–18^. Thus, elucidating how dietary ether lipids are incorporated into cells could enhance treatment strategies.

Ether lipids can be synthesized *de novo* from glycolytic intermediates or obtained from the diet. While the metabolic pathways involved in ether lipid synthesis are well-established, several key enzymes remain unidentified^19^. In some cases, enzymatic activities essential for ether lipid biosynthesis have been detected in tissue extracts, but the corresponding proteins have yet to be discovered^19^. These inferred enzymes without known sequences are referred to as orphan enzymes. Notably, the 1-O-alkyl-2-acetyl-*sn*-glycero-3-phosphate phosphatase (aka, alkylacetylglycerophosphatase) and the 1-O-alkylglycerol kinase are orphan enzymes in ether lipid biosynthesis (**Figure 1B, steps 5 and 8**).

Following the digestion of ether lipids, intestinal enzymes hydrolyze the *sn*-2 acyl group and the *sn*-3 phosphate-linked head group, but the ether bond remains resistant to degradation (**Figure 1B, step 7**). The resulting 1-O-alkylglycerols can be reincorporated into the host’s biomass, indicating the existence of an ether lipid salvage pathway^19^. To re-enter the biosynthetic pathway, 1-O-alkylglycerol undergoes phosphorylation by 1-O-alkylglycerol kinase, an orphan activity first detected over 50 years ago^20–22^ (**Figure 1B, step 8**). During the *de novo* synthesis of platelet activating factor, 1-O-alkyl-*sn*-glycero-3-phosphate is acetylated at the *sn*-2 position (**Figure 1B, step 4**), and the phosphate group is then removed by the orphan enzyme 1-O-alkyl-2-acetyl-*sn*-glycero-3-phosphate phosphatase (**Figure 1B, step 5**). This activity was first identified in 1986, with a noted preference for short-chain acyl groups at the *sn*-2 position of the glycerol backbone^23,24^. Here, we demonstrate that the kinase and phosphatase domains of ACAD10 function as the orphan 1-O-alkylglycerol kinase and the 1-O-alkyl-2-acetyl-*sn*-glycero-3-phosphate phosphatase, respectively.

## Results

### The ACAD10 kinase domain phosphorylates 1-O-alkylglycerols

While bioinformatically searching for divergent members of the protein kinase superfamily, we identified ACAD10, which contains a highly conserved, atypical kinase domain^25^. Additionally, ACAD10 includes haloacid dehalogenase-like hydrolase/phosphatase and long-chain acyl-CoA dehydrogenase domains, the latter of which has weak dehydrogenase activity toward 2-methyl-pentadecanoyl-CoA in vitro^25^. In higher eukaryotes, the kinase domain of ACAD10 is also present in its paralog, ACAD11. However, while both proteins share the dehydrogenase domain, ACAD11 lacks the phosphatase domain. ACAD10 is localized to peroxisomes, cytosol and mitochondria, whereas ACAD11 is found in the peroxisome^26,27^.

In *C. elegans*, the single ACAD10 gene (*acds-10*) encodes all three domains, whereas ACAD11 is absent (**Figure 1C**). To gain insight into the function of ACAD10, we purified the *C. elegans* ACAD10 kinase domain (residues 209-568, **Figure S1A**), incubated it with a crude mitochondrial extract from mouse liver and [γ-^32^P]-ATP, and then separated the reaction products by SDS-PAGE. We observed the incorporation of ^32^P into a low molecular weight species that was resistant to proteinase K treatment and required both acetyl-CoA and the active form of the *C. elegans* ACAD10 kinase domain for its formation (**Figure 1D**). Given that phospholipids migrate near the dye front during SDS-PAGE^28^, we hypothesized that ACAD10 might be phosphorylating a lipid. Therefore, we extracted the lipids from the reaction, separated them by thin layer chromatography (TLC), and observed two distinct ³²P-labelled species (**Figure 1E**). The reaction was time-dependent and required acetyl-CoA and Mg^2+^ as the activating divalent cation (**Figures 1E and S1B, C**). In addition to acetyl-CoA, CoA alone or other CoA thioesters also activated the *C. elegans* ACAD10 kinase domain (**Figure S1D**).

To identify the phosphorylated lipids, we treated the reaction products with strong base (0.5M KOH, saponification), extracted the lipids, and observed an increase in the ^32^P signal of one lipid species, accompanied by the disappearance of the other (**Figure 1F**). These results indicate that the *C. elegans* ACAD10 phosphorylates a non-saponifiable lipid, suggesting that ether lipids––known for their resistance to saponification––may be potential targets. Interestingly, the orphan 1-O-alkylglycerol kinase (**Figure 1B, step 8**) requires Mg^2+^ and acetyl-CoA^20–22^. Therefore, we hypothesized that ACAD10 may be the missing 1-O-alkylglycerol kinase in the ether lipid salvage pathway. Indeed, *C. elegans* ACAD10 phosphorylated chimyl alcohol (1-O-hexadecyl-*rac*-glycerol) (**Figure 1G**), as well as 1-O-acylglycerol (**Figure S1E**). Consistent with previous reports of 1-O-alkylglycerol kinase activity^21^, ACAD10 did not phosphorylate 1-O-alkyl-2-acyl-sn-glycerol (**Figure S1F**). Thus, ACAD10 phosphorylates both ether- and ester-linked glycerols in vitro but does not tolerate a long-chain acyl group at the sn-2 position.

### Structural and evolutionary insights into the ACAD10 kinase domain

The kinase domain of ACAD10 displays remarkable evolutionary conservation, with sequence identity reaching up to 63% between the human protein and its closest prokaryotic homologs (**Figure S2A**). We analyzed 2,101 organisms containing alkylglycerone phosphate synthase (AGPS)—the enzyme responsible for the first committed step in ether lipid biosynthesis—and found that 55% also encode the ACAD10 kinase domain (**Figure S2B**). In metazoans, 56% of species possess both AGPS and ACAD10, whereas fewer than 1% have ACAD10 alone, suggesting a strong selective pressure for their co-occurrence. We purified the kinase domains of several prokaryotic ACAD10 homologs from various species to assess their ability to phosphorylate 1-O-alkylglycerols **(Figure S2C)**. As expected, multiple homologs phosphorylated chimyl alcohol, with some showing acetyl-CoA dependence for full activity (**Figure S2D**).

We determined the crystal structure of the *Caldalkalibacillus thermarum* ACAD10 kinase domain bound to the ATP analog AMP-PNP at a resolution of 2.15 Å (**Figures 2A and 2B and Table S1**). *C. thermarum* ACAD10 exhibits a protein kinase/aminoglycoside phosphotransferase (APH) fold, characterized by a β-strand rich N-lobe and an α-helix-rich C-lobe. Structural similarity searches^29^ revealed that the APHs from *Cupriavidus pinatubonensis* (PDB: 3dxp) and *Mycobacterium tuberculosis* (PDB: 3ats) are the most structurally similar proteins (**Figure S2E-G**).

**Figure 2.**
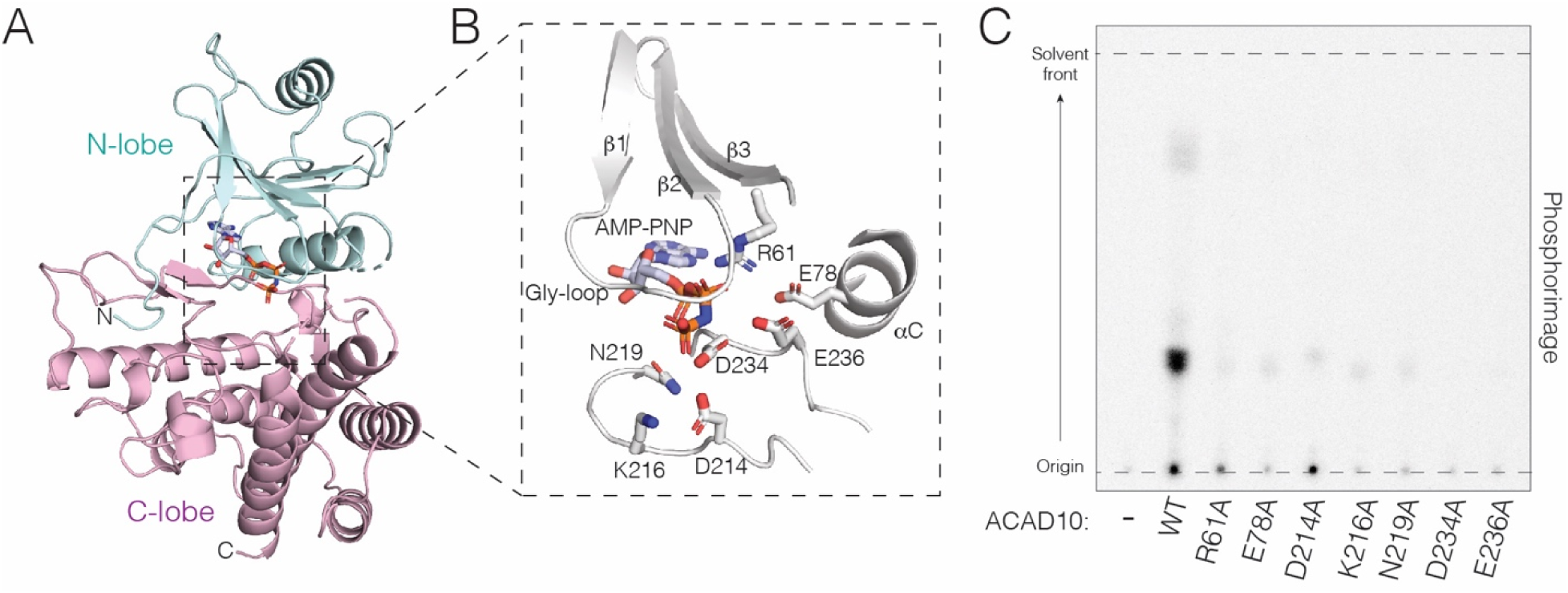
Structural insights into the ACAD10 kinase domain. **(A)** Cartoon representation of the *C. thermarum* ACAD10 kinase domain bound to the ATP analog AMP-PNP. The N-lobe and C-lobe are in light blue and pink, respectively. The AMP-PNP is in stick. **(B)** Zoomed in view of the *C. thermarum* ACAD10 kinase active site. AMP-PNP and the active site residues are shown as sticks. **(C)** Thin layer chromatogram depicting the incorporation of ^32^P from [γ-^32^P]ATP, into chimyl alcohol by *C. thermarum* ACAD10 or the active site mutants. Reactions were performed in the presence of acetyl-CoA and products were separated by TLC and visualized by phosphorimaging. Data are representative of at least three independent experiments.

Within the C-lobe lie several catalytic residues, including D214 (corresponding to protein kinase A; PKA: D166), N219 (PKA: N171), and D234 (PKA: D184). The nucleotide resides in a groove between the N and C lobes of the kinase domain. In canonical kinases, the lysine from the β3 strand forms an ion pair with the glutamate from the αC helix, indicating the active state of the kinase ^30^. However, in *C. thermarum* ACAD10, the β3 lysine is replaced with an arginine (R61), which is stabilized by the αC helix glutamate (E78) and interacts with the α-and β-phosphates of AMP-PNP (**Figure 2B**). Alanine substitution of R61 and E78 markedly reduced 1-O-alkylglycerol kinase activity (**Figure 2C**). In the C-lobe, the catalytic base D214 is situated within an HY<u>D</u>xK motif and is stabilized by the neighboring K216. Both residues are essential for *C. thermarum* ACAD10 activity (**Figure 2C**). In canonical kinases, this Asp is part of the HRD motif, which includes an Arg residue that binds to the phospho-amino acid group within the activation loop. However, both *C. thermarum* ACAD10 and eukaryotic ACAD10s lack activation loops, making them unlikely to be regulated by phosphorylation in this manner.

In *C. thermarum* ACAD10, the metal-binding residues N219 and D234 are located within the active site. Although Mg^2+^ was included in the crystallization conditions, we did not observe any electron density to model it into our structure. D234 is part of a DWE motif, which differs from the DFG motif typically found in canonical kinases. Substituting alanine for N219, D234, and E236 significantly reduced ether lipid kinase activity (**Figure 2C**).

Although acetyl-CoA stimulates *C. thermarum* ACAD10 kinase domain and was included in the crystallization conditions, we did not observe discernible electron density for it. We therefore generated an AlphaFold3 model of the *C. thermarum* ACAD10 kinase domain bound to ATP and acetyl-CoA. The model predicts a high-confidence acetyl-CoA-binding site (ipTM = 0.92) adjacent to the nucleotide-binding pocket, with the conserved DWE motif contacting two Mg^2+^ ions and the acetyl-CoA (**Figure S2H)**, providing a possible structural basis for acetyl-CoA-dependent activation. Collectively, our structural and AlphaFold3 analyses of the *C. thermarum* ACAD10 kinase domain identify several features that contribute to nucleotide binding, catalysis, and provide insight into acetyl-CoA-dependent activation.

### The ACAD10 phosphatase domain dephosphorylates 1-O-alkyl-2-acetyl-sn-glycero-3-phosphate

Platelet activating factor (PAF) is a potent bioactive ether lipid that drives platelet aggregation, vasodilation, inflammation, allergic responses, and shock ^31^. Structurally, PAF is unique among ether lipids due to the presence of an acetyl group at the *sn*-2 position of the glycerol backbone. It can be synthesized via two distinct pathways: the remodeling pathway (**Figure 3A**) or the *de novo* pathway (**Figure 3B**). While the remodeling pathway is the primary source of PAF under pathological conditions, the *de novo* pathway sustains basal PAF levels during normal cellular function ^32^. In the *de novo* pathway, an acetyl group is added to 1-O-alkyl-*sn*-glycero-3-phosphate, producing 1-O-alkyl-2-acetyl-*sn*-glycero-3-phosphate (**Figure 3B, step 1**). The phosphate is then removed by an orphan phosphatase that preferentially targets phospholipids containing short-chain acyl groups at the *sn*-2 position of the glycerol backbone. (**Figure 3B, step 2**) ^23,24^.

**Figure 3.**
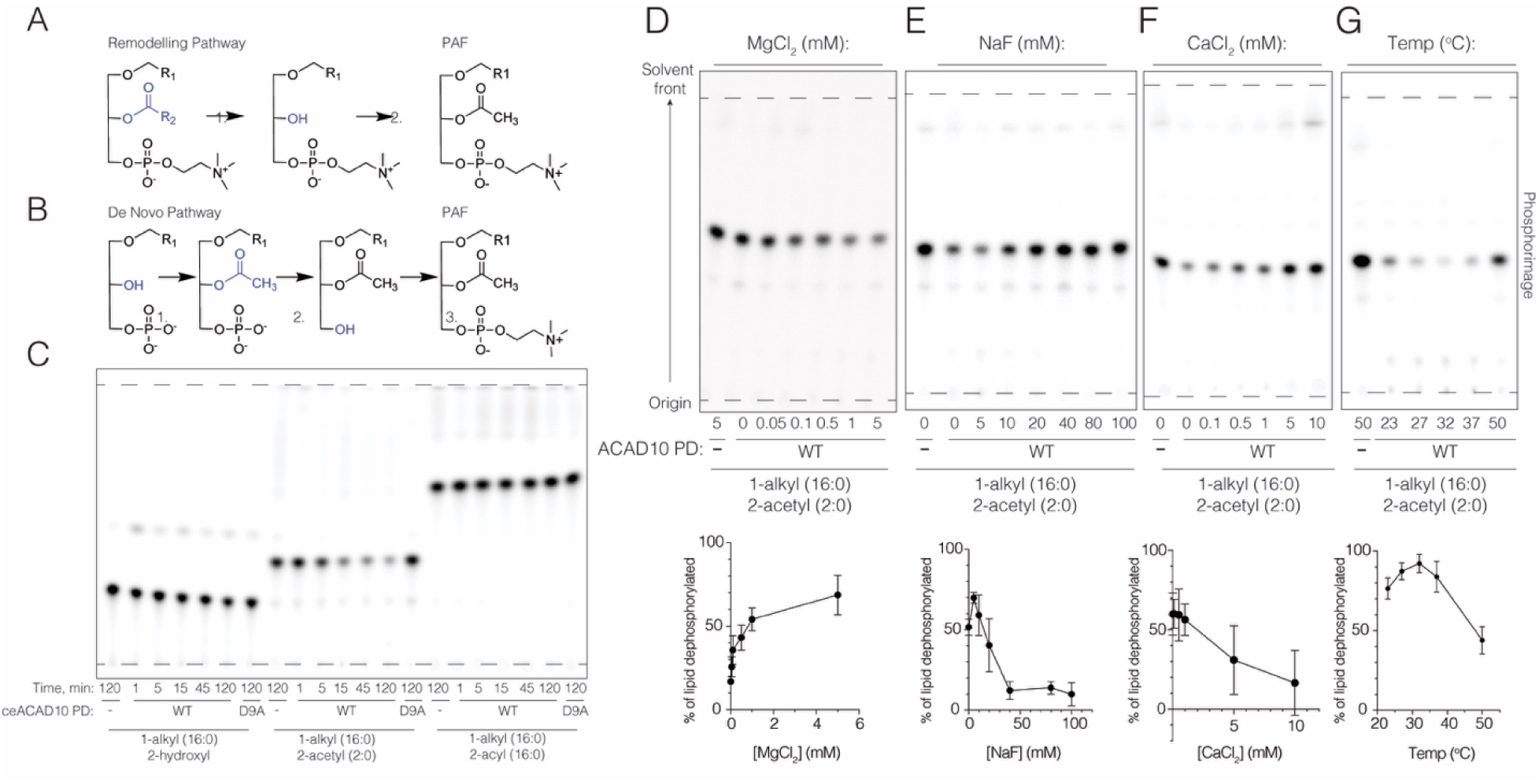
*C. elegans* ACAD10 phosphatase domain is a 1-O-alkyl-2-acetyl-*sn*glycero-3-phosphate phosphatase. **(A, B)** The remodeling (**A**) and *de novo* (**B**) pathways for PAF biosynthesis. In the remodeling pathway, a long-chain acyl group at the *sn*-2 position of the glycerol backbone is first removed by a phospholipase (**step 1**), followed by acetylation (**step 2**) to produce PAF. In the *de novo* pathway (see also steps **4-6** in Figure 1B), acetylation at the *sn*-2 position by AGPAT enzymes (**step 1**) is followed by the dephosphorylation of 1-O-alkyl-2-acetyl-*sn*-glycero-3-phosphate by an orphan phosphatase (**step 2**). Finally, a phosphatidylcholine (PC) headgroup is added in **step 3** to complete the synthesis of PAF. Adapted from^19^. **(D)** Thin layer chromatograms depicting the time-dependent dephosphorylation of ^32^P-labelled 1-O-alkyl-2-hydroxyl-*sn*-glycero-3-phosphate (1-alkyl (16:0), 2-hydroxyl), 1-O-alkyl-2-acetyl-*sn*-glycero-3-phosphate (1-alkyl (16:0), 2-acetyl (2:0)), and 1-O-alkyl-2-acyl-*sn*-glycero-3-phosphate (containing 16-carbon chain at *sn*-2; right) by *C. elegans* ACAD10 phosphatase domain or the inactive D9A mutant (DA). Reaction products were separated by TLC and visualized by phosphorimaging. Quantification of three independent experiments are shown in **Fig. S3B**. Data represent mean ± SD. **(E)** Thin layer chromatogram depicting ^32^P-labeled 1-O-alkyl-2-acetyl-*sn*-glycero-3-phosphate following treatment with the *C. elegans* ACAD10 phosphatase domain in the presence of increasing concentrations of MgCl_2_. Quantification of three independent experiments are shown below. Data represent mean ± SD. **(E-G)** Thin layer chromatograms depicting ^32^P-labeled 1-O-alkyl-2-acetyl-*sn*-glycero-3-phosphate following treatment with the *C. elegans* ACAD10 phosphatase domain in the presence of increasing concentrations of NaF (**E**), CaCl_2_ **(F)** and at different temperatures **(G)**. Reaction products were separated by TLC and visualized by phosphorimaging. Quantification of three independent experiments are shown below the respective chromatograms. Data represent mean ± SD.

To test whether ACAD10 encodes the missing 1-O-alkyl-2-acetyl-*sn*-glycero-3-phosphate phosphatase, we incubated the *C. elegans* ACAD10 phosphatase domain (residues 1-212, **Figure S3A**) with various ^32^P-labeled phospholipids, separated the reaction products by TLC, and detected ^32^P by phosphorimaging. The *C. elegans* ACAD10 phosphatase domain selectively dephosphorylated 1-O-alkyl-2-acetyl-*sn*-glycero-3-phosphate but not 1-O-alkyl-2-acyl-*sn*-glycero-3-phosphate, the latter of which contains a 16-carbon chain at the *sn*-2 position **(Figure 3C, S3B)**. The reaction was time-dependent and required Mg^2+^ as a cofactor **(Figure 3D)**. Notably, the ACAD10 phosphatase domain displayed several biochemical properties similar to those of the orphan phosphatase ^23,24^, including activity toward 1-O-alkyl-*sn*-glycero-3-phosphate **(Figure 3C, S3B)** and 1-O-acylglycerols with either a hydroxyl or an acetyl group at the *sn*-2 position of the glycerol backbone **(Figure S3C)**. Furthermore, like the orphan phosphatase, the ACAD10 phosphatase domain was inhibited by NaF **(Figure 3E)** and CaCl_2_ **(Figure 3F)** and displayed reduced activity at 23°C **(Figure 3G)**. Notably, substrate recognition by the ACAD10 phosphatase domain was dictated primarily by the substituent at the sn-2 position, with a strong preference for acetyl-containing substrates whereas both alkyl and acyl linkages at the sn-1 position were readily tolerated (**Figure S3B, C**), The predicted structure of the ACAD10 haloacid dehalogenase domain most closely resembles the lipid phosphatase domain of human bifunctional epoxide hydrolase 2 (EPHX2) and contains a five-helical cap predicted to contribute to the substrate-binding pocket (**Figure S3D**), potentially explaining its preference for acetyl-containing substrates. Thus, the *C. elegans* ACAD10 phosphatase domain has biochemical properties that closely resemble those of the orphan 1-O-alkyl-2-acetyl-*sn*-glycero-3-phosphate phosphatase.

### ACAD10-deficient *C. elegans* fail to incorporate dietary alkylglycerols into their cellular ether lipid pool

While mammalian ACAD10 contains a mitochondrial targeting sequence (MTS), *C. elegans* ACAD10 lacks the MTS but contains a predicted C-terminal peroxisomal targeting motif, S-R-L ^33^. When fused to an N-terminal mCherry tag, *C. elegans* ACAD10 localized to the peroxisome **(Figure 4A).** We generated ACAD10 knockout *C. elegans* (*acds-10-/-*) to investigate the role of ACAD10 in ether lipid biosynthesis. Compared to WT animals, *acds-10-/-* worms showed reduced fecundity and a shortened lifespan (**Figures 4B, C**). The observed reduction in lifespan aligns with a recent report indicating that ether lipid biosynthesis contributes to lifespan extension in *C. elegans* ^34^.

**Figure 4.**
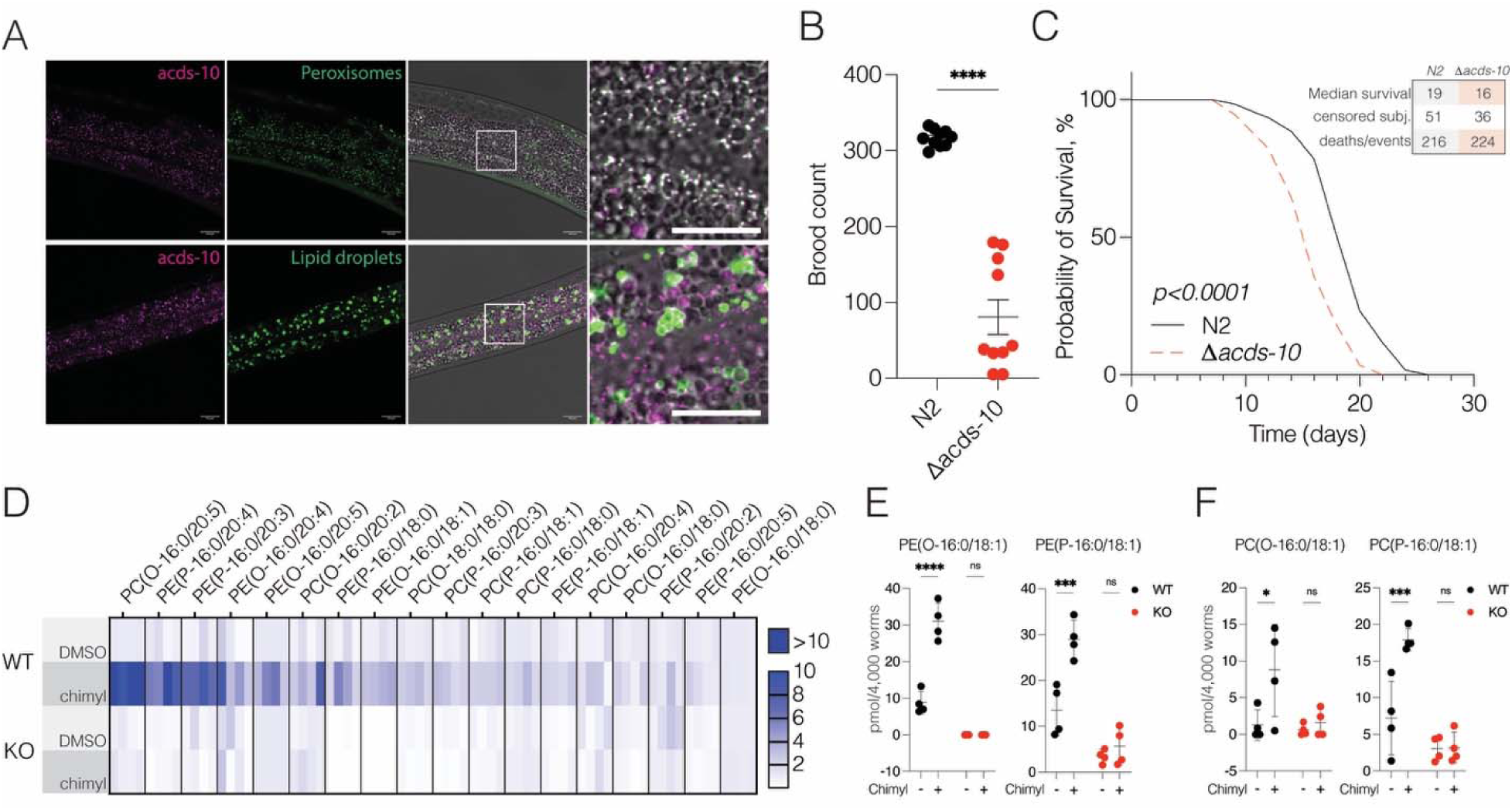
ACAD10 is required for proper ether lipid metabolism in *C. elegans*. **(A)** Fluorescent microscopy images of endogenous N-terminally mCherry-tagged *C. elegans* ACAD10 (magenta), the peroxisomal marker *3XFLAG::GFP::SKL* (green, upper) or the lipid droplet marker *DHS-3::GFP* (green, lower). Scale bar: 10 µm. **(B)** Graph illustrating progeny counts of control, N2 (WT), and *acds-10-/-* (KO) *C. elegans.* Data are presented as mean +/- SD. Each point represents the progeny count of an individual worm (n=10). **(C)** Kaplan–Meier curve depicting the survival of WT and *acds-10-/- C. elegans*. **(D)** Heatmap depicting the levels of 16:0 alkyl/plasmenyl chain ether lipid levels in WT and *acds-10-/- C. elegans* under normal dietary conditions and after dietary supplementation with chimyl alcohol. Each cell represents the fold change for an individual replicate of 4,000 worms, normalized to the average value of the unfed WT condition. **(E) F)** Representative ether lipids in WT and *acds-10-/- C. elegans* following dietary chimyl alcohol supplementation: PE(O-16:0/18:1) and PE(P-16:0/18:1) **(E)**; PC(O-16:0/18:1) and PC(P-16:0/18:1) **(F)**. Data represent mean ± SD, points represent biological replicates containing 4,000 worms, n = 4. Data was analyzed using two-way ANOVA with Šídák multiple comparisons test, ns: p>0.05, *p<0.05, ***p<0.001, ****p<0.0001.

*C. elegans* are typically cultured on the *E. coli* strain OP50, which lacks ether lipids^8^. Therefore, we supplemented their diets with *E. coli* enriched with chimyl alcohol (1-O-hexadecyl-*sn*-glycerol, O-16:0) and conducted lipidomics analysis to quantify lipid levels (**Figure S4A-G**). While WT animals fed chimyl alcohol showed an increase in phosphatidylcholine (PC)- and phosphatidylethanolamine (PE)-linked ether lipids, *acds-10-/-* animals largely lost the ability to salvage dietary chimyl alcohol (**Figures 4D-F and S4G-I**).

To identify the activities of *C. elegans* ACAD10 that are essential for these phenotypes, we generated strains with active site mutations in the phosphatase (D9A), kinase (D424A), and dehydrogenase (D966A) domains (**Figure 5A**) and monitored ether lipid levels after dietary supplementation of chimyl alcohol. While both the phosphatase and kinase activities were necessary for effective dietary ether lipid salvage, the dehydrogenase domain was largely dispensable (**Figures 5B**, **5C and S5**). Thus, the kinase and phosphatase activities of ACAD10 are necessary for proper ether lipid salvage and metabolism in *C. elegans*.

**Figure 5.**
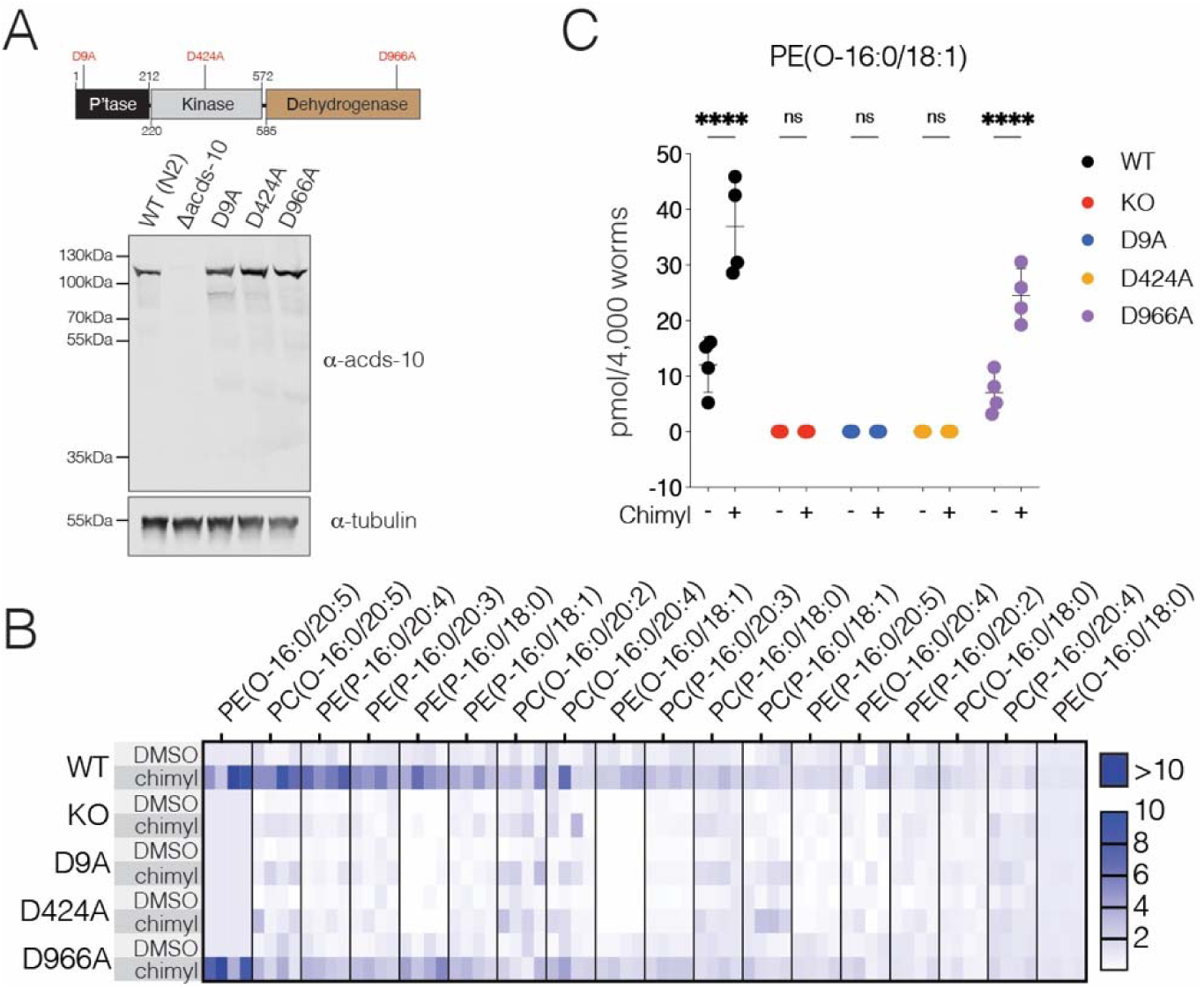
Both the kinase and phosphatase domains of ACAD10 are essential for proper ether lipid metabolism in *C. elegans*. **(A)** Protein immunoblots showing ACAD10 (acds-10) levels in *C. elegans* WT, *acds-10- /-,* and strains carrying the D9A, D424A, and D966A mutations. Tubulin is also shown as a loading control. Domain architecture of *C. elegans* ACAD10, highlighting the locations of these mutations, is displayed above the immunoblot. **(B)** Heatmap depicting the levels of 16:0 alkyl/plasmenyl chain ether lipid levels in WT, *acds-10-/-*, and *C. elegans* strains harboring mutations in the phosphatase (D9A), kinase (D424A), and dehydrogenase (D966A) domains under normal dietary conditions and after dietary supplementation with chimyl alcohol. Each cell represents the fold change for an individual replicate of 4,000 worms, normalized to the average value of the unfed WT condition. Data represent mean ± SD, n=4. **(C)** Levels of a representative ether lipid, PE(O-16:0/18:1), in WT, *acds-10-/-*, and mutant *C. elegans* strains D9A, D424A, and D966A under normal dietary conditions (-) and following chimyl alcohol supplementation (+). Data represent mean ± SD, points represent biological replicates containing 4,000 worms, n=4. Data was analyzed using two-way ANOVA with Šídák multiple comparisons test, ns: p>0.05, ****p<0.0001.

### ACAD10 is essential for the incorporation of dietary alkylglycerols into the ether lipid pool in mice

Recently, ACAD10 and ACAD11 were shown to phosphorylate 4-hydroxyacyl-CoAs to form 4-phosphoacyl-CoAs, which are then converted by the dehydrogenase domains to 2-enoyl-CoAs in vitro and in mammalian cells^26,35^. Plasma levels of 4-hydroxy acids were elevated in Acad11 knockout mice^26^. However, 4-hydroxy acid levels in Acad10 KO mice have not been evaluated. To determine whether mammalian Acad10 is involved in the metabolism of 4-hydroxy acids or ether lipids *in vivo*, we assessed 4-hydroxy acids under basal conditions and ether lipids following dietary supplementation with batyl alcohol (1-O-octadecyl-rac-glycerol, O-18:0) in Acad10 KO mice (**Figure S6A-E**). Through a targeted analysis of oxidized lipids in the plasma of Acad10 KO mice, we observed a reduction in inflammatory oxylipins (**Figures S6F and S6G**). However, levels of 4-hydroxy acids remained unchanged between control and Acad10 KO mice (**Figures 6A and S6H-I**). Additionally, *acds-10-*deficient *C. elegans* exhibited no changes in 4-hydroxy acid levels (**Figure 6B**).

**Figure 6.**
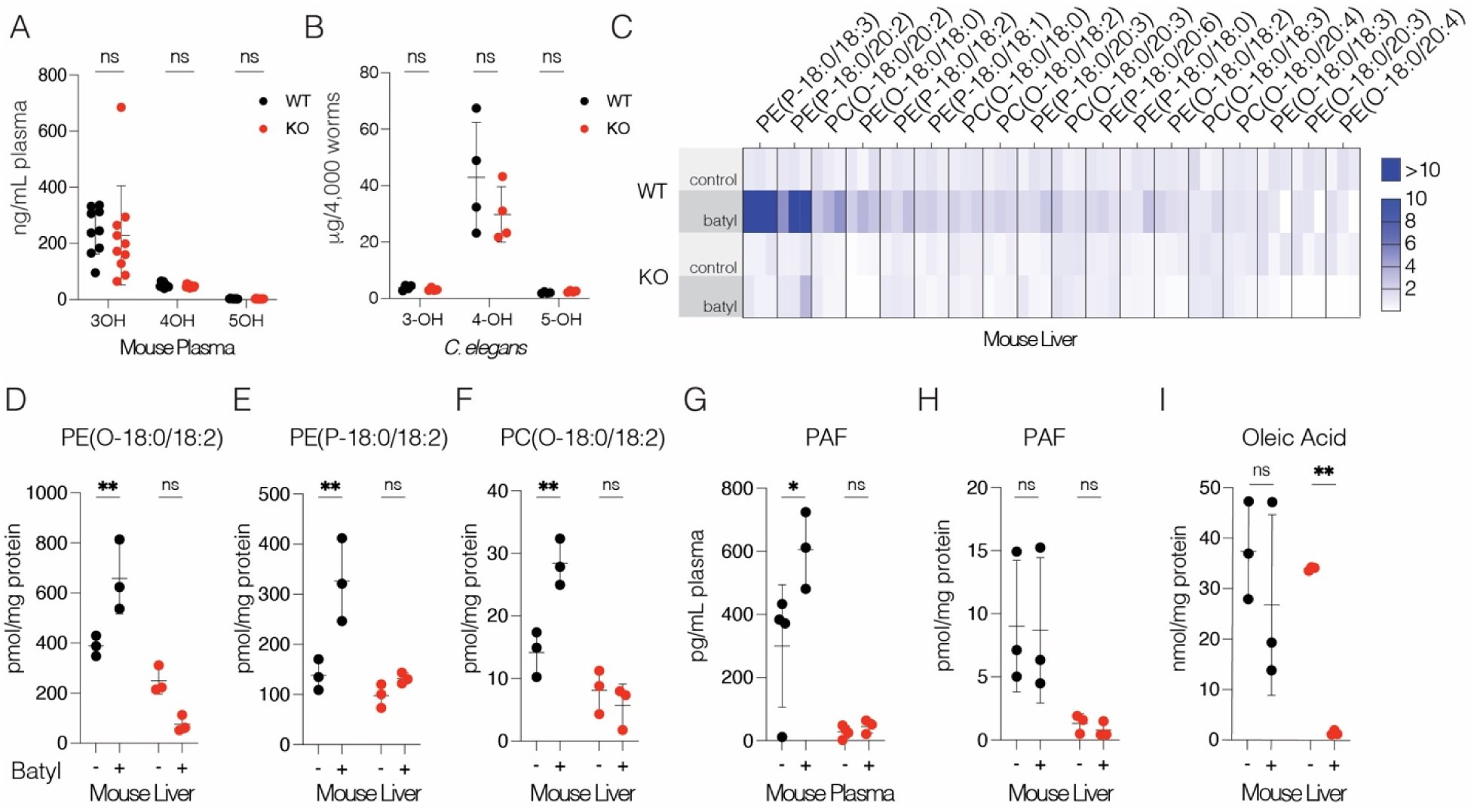
ACAD10 is essential for proper ether lipid metabolism in mice. **(A)** Plot showing combined plasma levels of 3-, 4-, and 5-hydroxy fatty acid species in WT and Acad10 KO mice. Data represent mean ± SD, points represent individual mice, n=9 and n=10 for WT and Acad10 KO mice, respectively. Data was analyzed using two-tailed unpaired t-tests for each hydroxy fatty acid group, ns: p>0.05. **(B)** Plot showing combined levels of 3-, 4-, and 5-hydroxy fatty acid species in WT and *acds-10*-/- *C. elegans*. Data represent mean ± SD, points represent biological replicates containing 4,000 worms, n=4. Data was analyzed using two-tailed unpaired t-tests for each hydroxy fatty acid group, ns: p>0.05. **(C)** Heatmap showing liver 18:0 alkyl/plasmenyl chain ether lipid levels in WT and Acad10 KO mice under normal dietary conditions and after batyl alcohol supplementation. Each cell represents the fold change for an individual mouse, normalized to the average value of the unfed WT condition. **(D-F)** Representative liver ether lipids in WT and Acad10 KO mice following dietary batyl alcohol supplementation (+): PE(O-18:0/18:2) **(D),** PE(P-18:0/18:2) **(E)**; and PC(O-18:0/18:2) **(F)**. Data represent mean ± SD, points represent individual mice, n = 3 mice per group. Data was analyzed using two-way ANOVA with Šídák multiple comparisons test, ns: p>0.05, **p<0.01. **(G-H)** Plasma **(G)** and liver **(H)** PAF levels in WT and Acad10 KO mice after dietary batyl alcohol supplementation (+). Data represent mean ± SD, points represent individual mice, n=3 mice per group. Data was analyzed using two-way ANOVA with Šídák multiple comparisons test, ns: p>0.05, *p<0.05. **(I)** Liver oleic acid (18:1) levels of WT and Acad10 KO mice following dietary batyl alcohol supplementation (+). Data represent mean ± SD, points represent individual mice, n = 3 mice per group. Data was analyzed using two-way ANOVA with Šídák multiple comparisons test, ns: p>0.05, **p<0.01.

Consistent with our findings in worms, dietary supplementation with batyl alcohol elevated ether lipid levels in the livers of WT mice, but not of Acad10 KO mice **(Figures 6C-F and S6J-L).** Notably, in the absence of batyl alcohol supplementation, ether lipid levels in the livers of Acad10 knockout mice remained largely unchanged, whereas liver and plasma PAF levels were reduced (**Figures 6G, H and S6M-P**). These findings are consistent with the involvement of the ACAD10 kinase domain in ether lipid salvage, and of the phosphatase domain in *de novo* PAF biosynthesis. While most other liver lipids remained unchanged (**Figures S6Q-V**), we observed some changes in free fatty acids, including a notable decrease in oleic acid levels only in Acad10 KO mice fed batyl alcohol (**Figures 6I and S6W)**. Collectively, these results indicate that ACAD10 plays a role in ether lipid biosynthesis and salvage in both worms and mice. Further, our results suggest that ACAD10 and ACAD11 may have distinct functions.

### Ether lipid kinase and phosphatase activities are conserved in human ACAD10 and ACAD11

Having established the biochemical activities of the isolated *C. elegans* ACAD10 kinase and phosphatase domains, we next asked whether these activities are conserved in the human ACAD10 and ACAD11 and how the opposing activities of ACAD10 are coordinated within a single polypeptide. AlphaFold3 models of full-length human ACAD10 did not predict stable interdomain interactions (**Figure 7A**). Instead, the catalytic domains appear to be connected by short flexible linkers. Recombinant human ACAD10 (residues 41–617), containing the phosphatase and kinase domains, and the ACAD11 kinase domain (residues 11–356) both phosphorylated 1-O-alkylglycerol and 1-O-alkyl-2-acetyl-*sn*-glycerol, but not 1-O-alkyl-2-acyl-*sn*-glycerol containing a long-chain acyl group at the sn-2 position (**Figures 7B and S7A, B**). Inactivation of the ACAD10 phosphatase domain markedly increased kinase product accumulation, demonstrating that the phosphatase reverses the kinase reaction in vitro. However, the 2-hydroxyl-containing kinase product was a much poorer phosphatase substrate than the corresponding 2-acetyl intermediate (**Figure 7C**), as observed for the isolated *C. elegans* phosphatase domain (**Figures 3C and S3B, C**). These findings suggest that substrate discrimination at the sn-2 position minimizes futile cycling between the opposing kinase and phosphatase activities.

**Figure 7.**
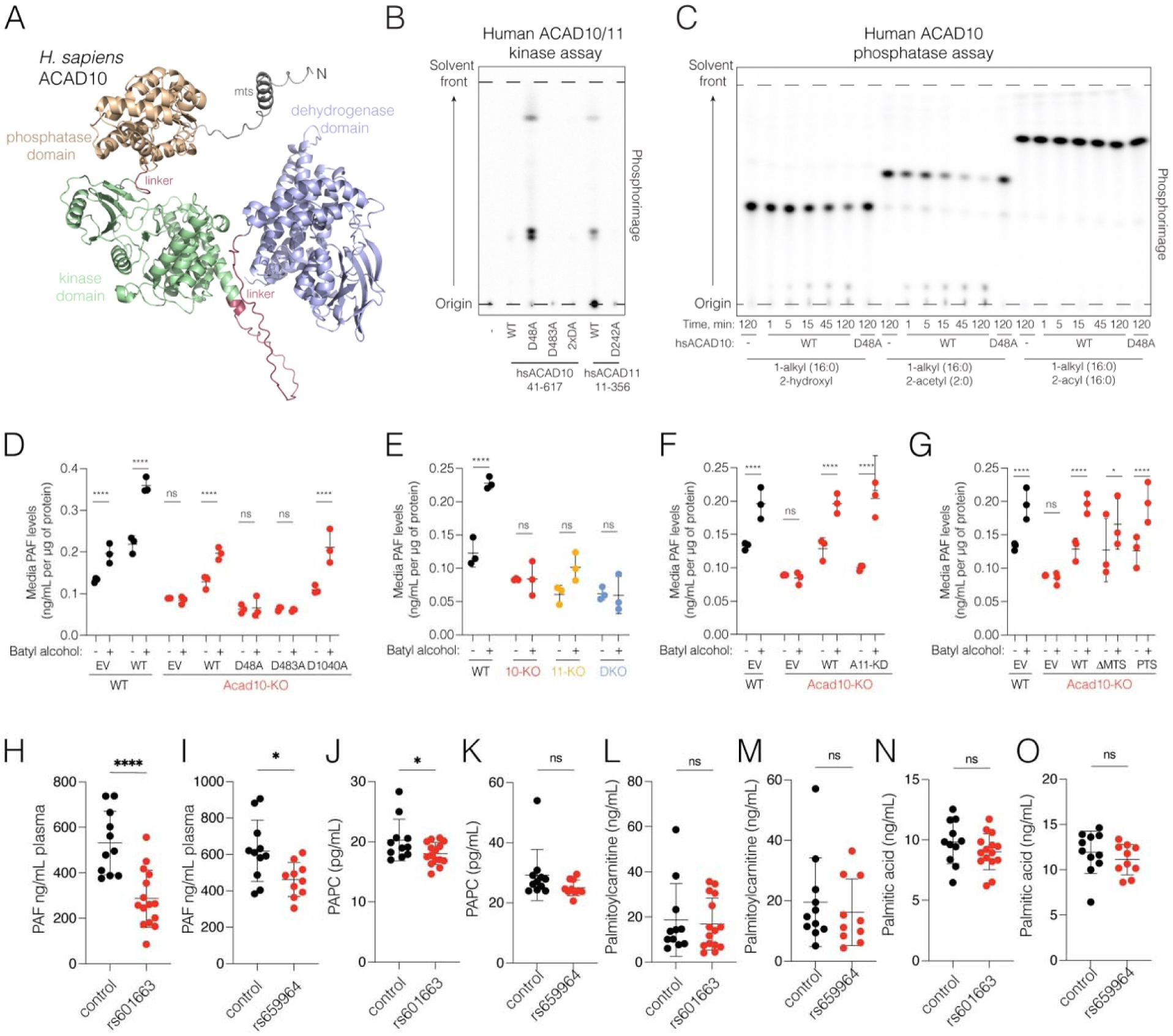
Human ACAD10 and ACAD11 regulate ether lipid metabolism. **(A)** AlphaFold3 model of human ACAD10. **(B)** Thin layer chromatogram depicting the incorporation of ^32^P from [γ-^32^P]ATP into chimyl alcohol by *H. sapiens* ACAD10 (residues 41–617), which contains the phosphatase (PD) and kinase (KD) domains, and the *H. sapiens* ACAD11 KD (residues 11–356). In ACAD10, D48A and D483A inactivate the PD and KD, respectively, and 2XDA denotes the D48A/D483A double mutant. In ACAD11, D242A inactivates the KD. Reaction products were separated by TLC and visualized by phosphorimaging. Data are representative of at least three independent experiments. **(C)** Thin layer chromatograms depicting the time-dependent dephosphorylation of ^32^P-labeled ether lipids containing different substituents at the sn-2 position by the *H. sapiens* ACAD10 PD or the inactive D48A mutant. Reaction products were separated by TLC and visualized by phosphorimaging. Data are representative of at least three independent experiments. **(D)** Media PAF levels in WT, Acad10-KO, and Acad10-KO Hepa1-6 cells expressing empty vector (EV), WT human ACAD10 (WT), or ACAD10 mutants that inactivate the phosphatase (D48A), kinase (D483A), or dehydrogenase (D1040A) domains. Cells were treated with 20 µM batyl alcohol for 24 h (+) or DMSO vehicle control (-). Data represent mean ± SD, with individual data points shown (n = 3). Data were analyzed using two-way ANOVA with Šídák multiple comparisons test; ns, p > 0.05; ****p < 0.0001. **(E)** Media PAF levels in WT, Acad10-KO (10-KO), Acad11-KO (11-KO), and Acad10/Acad11 double-KO (DKO) Hepa1-6 cells treated with 20 ∝M batyl alcohol for 24 h (+) or DMSO vehicle control (-). Data represent mean ± SD, with individual data points shown (n = 3). Data were analyzed using two-way ANOVA with Šídák multiple comparisons test; ns, p > 0.05; ****p < 0.0001. **(F)** Media PAF levels in WT, Acad10-KO, and Acad10-KO Hepa1-6 cells expressing empty vector (EV), WT human ACAD10 (WT), or an ACAD10 chimera in which the kinase domain was replaced with the human ACAD11 kinase domain (A11-KD). Cells were treated with 20 ∝M batyl alcohol for 24 h (+) or DMSO vehicle control (-). Data represent mean ± SD, with individual data points shown (n = 3). Data were analyzed using two-way ANOVA with Šídák multiple comparisons test; ns, p > 0.05; ****p < 0.0001. **(G)** Media PAF levels in WT, Acad10-KO, and Acad10-KO Hepa1-6 cells expressing empty vector (EV), WT human ACAD10 (WT), ACAD10 lacking the mitochondrial targeting sequence (ΔMTS; residues 41–C), or ΔMTS ACAD10 fused to the ACAD11 C-terminal peroxisomal targeting sequence (PTS). Cells were treated with 20 ∝M batyl alcohol for 24 h (+) or DMSO vehicle control (-). Data represent mean ± SD, with individual data points shown (n = 3). Data were analyzed using two-way ANOVA with Šídák multiple comparisons test; ns, p > 0.05; *p < 0.05; ***p < 0.001; ****p < 0.0001. **(H–O)** Plasma PAF **(H, I)**, 1-palmitoyl-2-acetyl-sn-glycero-3-PC (PAPC) **(J, K)**, palmitoylcarnitine **(L, M)**, and palmitic acid **(N, O)** levels in Akimel O’odham individuals carrying the rs601663 or rs659964 polymorphism and matched controls. Data represent mean ± SD, with individual participants shown; rs601663, n = 15; rs659964, n = 10; matched controls, n = 11. Data were analyzed using two-tailed unpaired t-tests; ns, p > 0.05; *p < 0.05; ****p < 0.0001.

Consistent with our worm and mouse studies, batyl alcohol increased PAF levels in WT but not *Acad10*-KO mouse hepatoma Hepa1-6 cells (**Figure 7D**). Re-expression of WT human ACAD10 or the dehydrogenase mutant (D1040A), but not phosphatase-inactive (D48A) or kinase-inactive (D483A) mutants, restored PAF levels, indicating that ether lipid salvage requires the kinase and phosphatase, but not dehydrogenase, activity of human ACAD10 (**Figure 7D and S7C**). Acad11-KO and ACAD10/11 double-KO (DKO) cells also displayed reduced ether lipid levels and were unresponsive to batyl alcohol (**Figure 7E**), indicating that both ACAD10 and ACAD11 contribute to ether lipid metabolism. Consistent with overlapping catalytic functions, an ACAD10 chimera containing the ACAD11 kinase domain restored PAF production in Acad10-KO cells (**Figure 7F**). Together, these data support overlapping catalytic functions for ACAD10 and ACAD11 but do not establish functional redundancy in vivo.

Although human ACAD10 contains an N-terminal MTS, it has been reported to localize to multiple subcellular compartments^25,27,36^. Interpretation of these localization studies may be complicated by a recent report that ACAD10 undergoes proteolytic cleavage between the kinase and dehydrogenase domains, generating distinct N- and C-terminal fragments^35^. We therefore tested whether the MTS is required for ACAD10-dependent ether lipid salvage. Deletion of the MTS (ACAD10 41–1059, ΔMTS) or addition of the ACAD11 C-terminal peroxisomal targeting sequence (PTS) to ΔMTS ACAD10 (ACAD10 1–1059 + ACAD11 771–780, PTS) restored ether lipid production and PAF levels as efficiently as full-length ACAD10 in Acad10-KO cells (**Figures 7G**). Thus, the MTS is dispensable for ACAD10-dependent ether lipid salvage in this cellular system.

### Human individuals with *ACAD10* polymorphisms have decreased plasma ether lipid levels

Genome-wide association studies (GWAS) in the Akimel O’odham tribe (Pima Indian Cohort Study) revealed a link between single nucleotide polymorphisms in the *ACAD10* gene and early-onset type 2 diabetes, diabetic kidney disease, and increased adiposity^37,38^. These metabolic dysfunctions are correlated with two polymorphisms in *ACAD10*, rs601663 (within the promoter) and rs659964 (within an intron)^37^. We collected plasma from 15 individuals heterozygous for the rs601663 variant with 11 matched controls, and 10 individuals heterozygous for the rs659964 variant with 11 matched controls to monitor plasma lipid levels (**Figure 7H-O**). Remarkably, individuals carrying the rs601663 and rs659964 variants exhibited a 46% and 25% reduction in PAF levels, respectively with minimal changes in the other lipids analyzed. (**Figures 7H, I**). Thus, ACAD10 is required for proper ether lipid metabolism in worms, mice and humans.

## Discussion

ACAD10 and ACAD11 have been linked to various metabolic disorders in both mice and humans^37,39,40^. GWAS across diverse human populations have identified polymorphisms in *ACAD10* and *ACAD11* that are associated with kidney and cardiovascular diseases^41,42^. Notably, *ACAD10* polymorphisms are also connected to type 2 diabetes in the Akimel O’odham tribe, a population with the highest recorded rates of type 2 diabetes (50%) and diabetic kidney disease (30%)^37,43,44^. Although PAF levels are reduced in individuals with the rs601663 and rs659964 variants, it remains uncertain how these reductions, along with the presumed decrease in dietary alkylglycerol salvage, contribute to diabetes and diabetic kidney disease. In mice with a mixed background (SvEv129/BL6), loss of ACAD10 leads to increased adiposity, elevated insulin levels, glucose intolerance, and impairments in the insulin signaling pathway^43^. However, a recent study found no metabolic phenotypes associated with ACAD10 loss in mice on a pure C57Bl/6J background^45^. The reason for this discrepancy is unclear, but variations in dietary challenge and genetic background between the studies may help explain the phenotypic differences. Based on our results linking ACAD10 to ether lipid biosynthesis, further studies will be necessary to evaluate potential metabolic phenotypes resulting from dietary ether lipid challenges.

The kinase and dehydrogenase domains of ACAD10 show sequence similarity to *Pseudomonas putida* enzymes LvaA and LvaC, which participate in levulinic acid catabolism through a 4-hydroxy acyl-CoA intermediate^26^. In bacteria, levulinic acid is a dehydration product of plant biomass and can serve as the sole carbon source for certain species^46^. In humans, 4-hydroxy acids are generated through lipid peroxidation, the breakdown of longer-chain hydroxy acids, or by the ingestion of certain drugs of abuse^26^. Recently, ACAD10 and ACAD11 were shown to phosphorylate 4-hydroxyacyl-CoA to form 4-phosphoacyl-CoA, which is then converted by the dehydrogenase domains to 2-enoyl-CoA ^26,35^. Although we did not observe any differences in 4-hydroxy acid levels in ACAD10 KO worms or mice, Rashan et al^26^. reported elevated plasma levels of 4-hydroxy acids in ACAD11 knockout mice. However, neither our study nor the referenced study tested ACAD10 or ACAD11 KO mice under conditions that enhance 4-hydroxy acid production. In any event, our results suggest that ACAD10 and ACAD11 may have distinct functions, or that they participates in multiple metabolic pathways. Further research is needed to determine whether Acad11 KO mice display alterations in ether lipid biosynthesis or salvage.

*C. elegans* ACAD10 localizes to peroxisomes (**Figure 4A**), whereas mammalian ACAD10 and ACAD11 have distinct subcellular localizations—mitochondria and peroxisomes, respectively^26^—suggesting divergent cellular functions. Although human ACAD10 contains an N-terminal MTS, it has also been reported to localize to multiple subcellular compartments^25,27,36^. Given that most ether lipid biosynthetic enzymes are associated with the ER or peroxisomes, the mitochondrial localization of mammalian ACAD10 is intriguing. Notably, when supplied exogenously to mammalian cells, ether lipids preferentially accumulate in mitochondria^47^, supporting our model that the kinase domain of ACAD10 serves as the initiating enzyme in the ether lipid salvage pathway.

Our findings in *C. elegans* suggest that the ether lipid-related activity of ACAD10 may represent the ancestral function of Metazoan ACAD10/11. While a previous report claimed that *C. elegans* ACAD10 mediates metformin’s anti-aging and anti-cancer effects^48^, the gene the authors studied, *F37H8.3*, is not a homolog of human ACAD10, contrary to their claims. *F37H8.3* encodes a phosphatase-like domain that is only distantly related to the phosphatase domain of human ACAD10 and lacks both the kinase and dehydrogenase domains.

In summary, we have identified two previously unknown enzymes involved in the biosynthesis and salvage of ether lipids. This discovery is expected to catalyze further research into the impact of dietary ether lipids on human health and disease. Given that the role of ether lipids in human biology remains largely unexplored, our findings may offer valuable insights into their biological functions.

### Limitations of the study

Although our findings establish ACAD10 as a key enzyme in ether lipid metabolism, its kinase and phosphatase domains can also act on acyl-containing lipid substrates in vitro. Whether these activities contribute to acyl lipid metabolism in vivo remains unknown. Moreover, while Acad11-KO Hepa1-6 cells display defects in ether lipid metabolism, the contribution of ACAD11 to ether lipid salvage and PAF production in vivo remains to be determined. The relationship between the previously reported 4-hydroxy fatty acid kinase activity of ACAD10/11^26,35^ and the ether lipid kinase and phosphatase activities described here also remains unclear.

## Supporting information

Manuscript File

## Resource availability

## Lead contact

Further information and requests for resources and reagents should be directed to and will be fulfilled by the lead contact, Vincent S. Tagliabracci

## Materials availability

All unique/stable reagents generated in this study are available from the lead contact with a completed Materials Transfer Agreement.

## Acknowledgements

We thank members of the Tagliabracci laboratory for helpful discussions. Results shown in this report are derived from work performed at the Argonne National Laboratory, Structural Biology Center at the Advanced Photon Source. SBC-CAT is operated by UChicago Argonne, LLC, for the US Department of Energy, Office of Biological and Environmental Research under contract DE-AC02-06CH11357. The contents of this publication are solely the responsibility of the authors and do not necessarily represent the official views of NIGMS or NIH.

This work was funded by NIH Grants DP2GM137419, R35GM158265 (V.S.T.), R01DK137976 (D.J.P), Welch Foundation Grants I-1911 (V.S.T.), the Howard Hughes Medical Institute (HHMI, V.S.T, J.S.), the Glenn Foundation and American Federation for Aging Research (A22068 to J.S.); Hatch Grant (WIS04000-1024796 to J.S. and R.J.); JDRF (JDRF201309442 to J.S.); an R01 through NIH/NIDDK (R01DK133479 to J.S.); Biotechnology Program T32 (5T32GM135066-05;TK), and support of the Biology of Aging and Age Related Diseases T32 (AG000213; D.A.B.). J.S is an HHMI Freeman Hrabowski Scholar. V.S.T. is a Michael L. Rosenberg Scholar in Medical Research, a CPRIT Scholar (RR150033), a Searle Scholar and an investigator of the HHMI.

## Author contributions

J.S.Y., E.P., V.A.L., L.T., P.M.D., J.S. and V.S.T. designed the experiments. K.P. performed the bioinformatics. J.S.Y., E.P., V.A.L., and V.S.T., performed molecular cloning, protein production and biochemical assays. J.S.Y., L.T., J.K. and P.M.D. performed *C. elegans* experiments. J.S.Y. and D.R.T. performed crystallization and structure determination. J.M., T.K., D.B., E.R., E.P., D.J.P., and J.S. performed mouse work, and lipidomics analyses. J.S.Y., E.P., P.M.D., and V.S.T. wrote the manuscript with input from all authors.

