## Supplementary material for "ACAD10 encodes two orphan enzymes in the ether lipid biosynthetic and salvage pathways": Manuscript File

**Supplemental Figures**

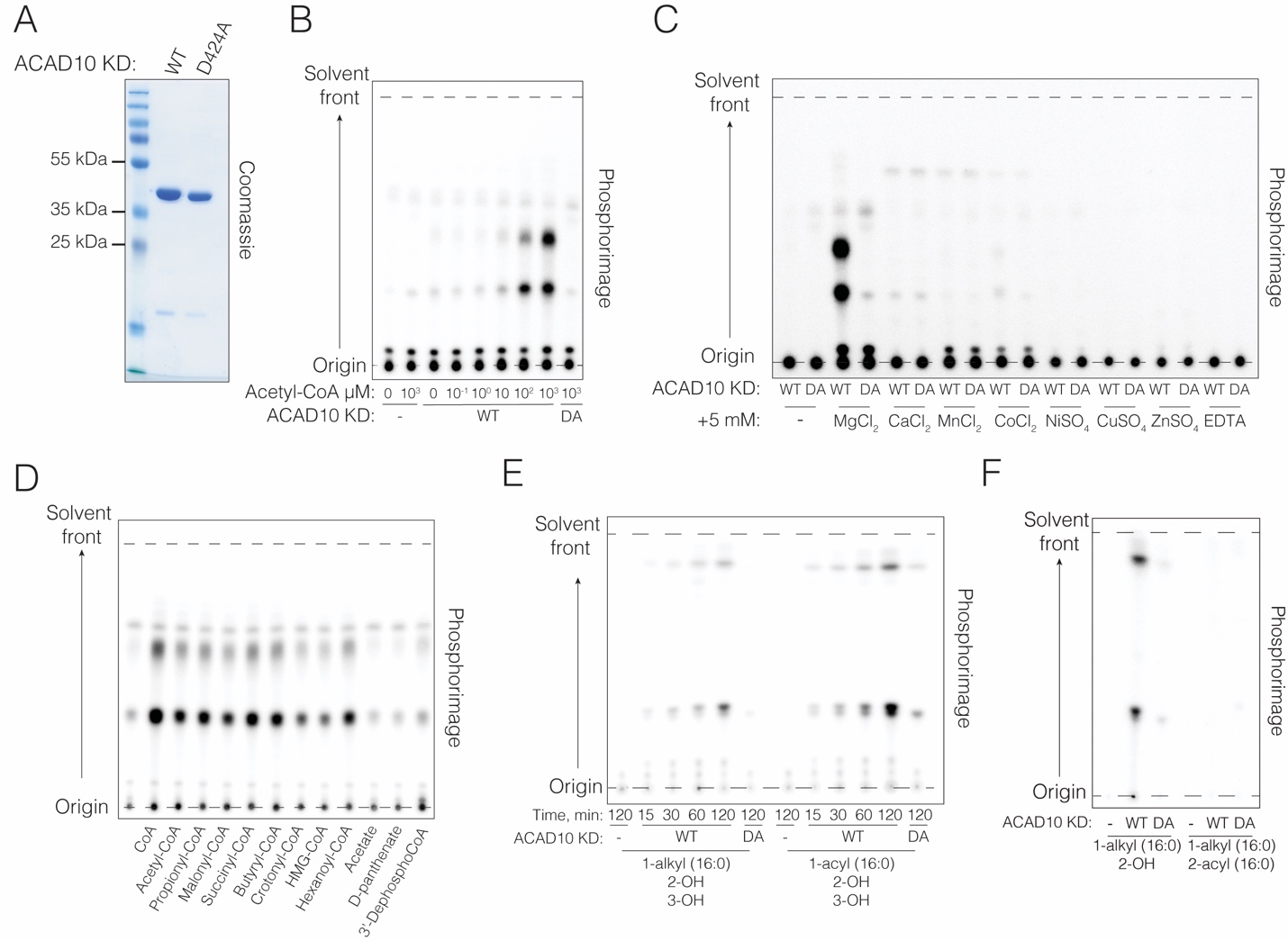

**Figure S1: Characterization of *C. elegans* ACAD10 kinase domain, related to Figure 1.**

**(A)** SDS-PAGE and Coomassie staining of recombinant *C. elegans* ACAD10 kinase domain and the D424A mutant.

**(B-D)** Thin layer chromatograms depicting the incorporation of ^32^P from [γ-^32^P]ATP into two lipid species from mouse liver mitochondrial extracts by *C. elegans* ACAD10 KD. Reactions were performed in the presence of increasing concentrations of acetyl-CoA **(B),** different divalent cations **(C)** and different CoA thioesters **(D)**. Lipids were extracted, separated by TLC and visualized by phosphorimaging. Data are representative of at least three independent experiments.

**(E)** Thin layer chromatogram depicting the incorporation of ^32^P from [γ-^32^P]ATP into ether (left) or acyl (right) lipid substrates by the *C. elegans* ACAD10 kinase domain. Reaction products were separated by TLC and visualized by phosphorimaging. Data are representative of at least three independent experiments.

**(F)** Thin layer chromatogram depicting the incorporation of ^32^P from [γ-^32^P]ATP into the 1-alkylglycerol chimyl alcohol, or 1-alkyl-2-acyl-sn-glycerol by *C. elegans* ACAD10 KD. Reaction products were separated by TLC and visualized by phosphorimaging. Data are representative of at least three independent experiments.

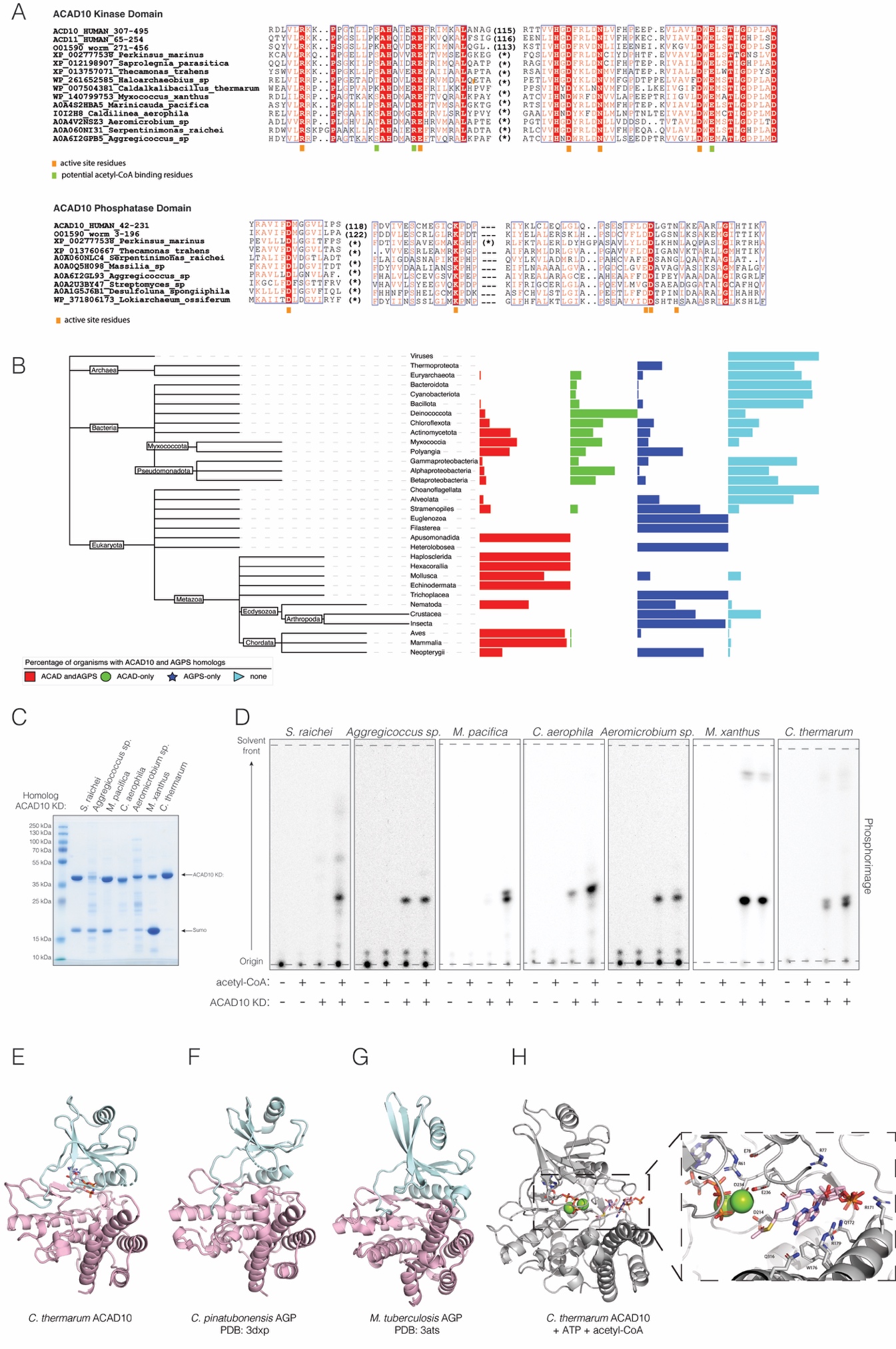

**Figure S2: Evolutionary and structural insights into the ACAD10 kinase domain, related to Figure 2.**

**(A)** Multiple sequence alignment highlighting conserved active-site residues in the ACAD10 kinase (top) and phosphatase (bottom) domains.

**(B)** Taxonomy dendrogram for selected taxa with histogram depicting percentages of organisms within a taxon possessing ACAD10 and AGPS homologs (red), ACAD10 homologs only (green), AGPS homologs only (dark blue) and neither (cyan). 22509 organisms from Uniprot Representative Proteomes analyzed.

**(C)** SDS-PAGE and Coomassie staining of recombinant ACAD10 kinase domains from various bacterial species.

**(D)** Thin layer chromatograms depicting the incorporation of ^32^P, from [γ-^32^P]ATP, into chimyl alcohol by bacterial *Serpentinimonas raichei*, *Aggregicoccus sp*., *Marinicauda pacifica*, *Caldilinea aerophile*, *Aeromicrobium sp,* *Myxococcus xanthus* and *Caldalkalibacillus thermarum* ACAD10 KDs. Reactions were performed in the presence or absence of acetyl-CoA and the reaction products were separated by TLC and visualized by phosphorimaging. Data are representative of at least three independent experiments.

**(E–G)** Cartoon representations of the *C. thermarum* ACAD10 kinase domain **(E)** and the aminoglycoside phosphotransferases (APHs) from *Cupriavidus pinatubonensis* (**F**; PDB: 3DXP) and *Mycobacterium tuberculosis* (**G**; PDB: 3ATS). The N- and C-lobes are colored light blue and pink, respectively.

**(H)** AlphaFold model of the *C. thermarum* ACAD10 kinase domain (KD; left) and a magnified view of the active site (right), showing ATP, and acetyl-CoA rendered as sticks. Predicted interaction confidence scores (chain pair ipTM) were 0.91 for ACAD10-KD–acetyl-CoA, 0.95 for ACAD10-KD–ATP, and 0.95 for ACAD10-KD–Mg^2+^.

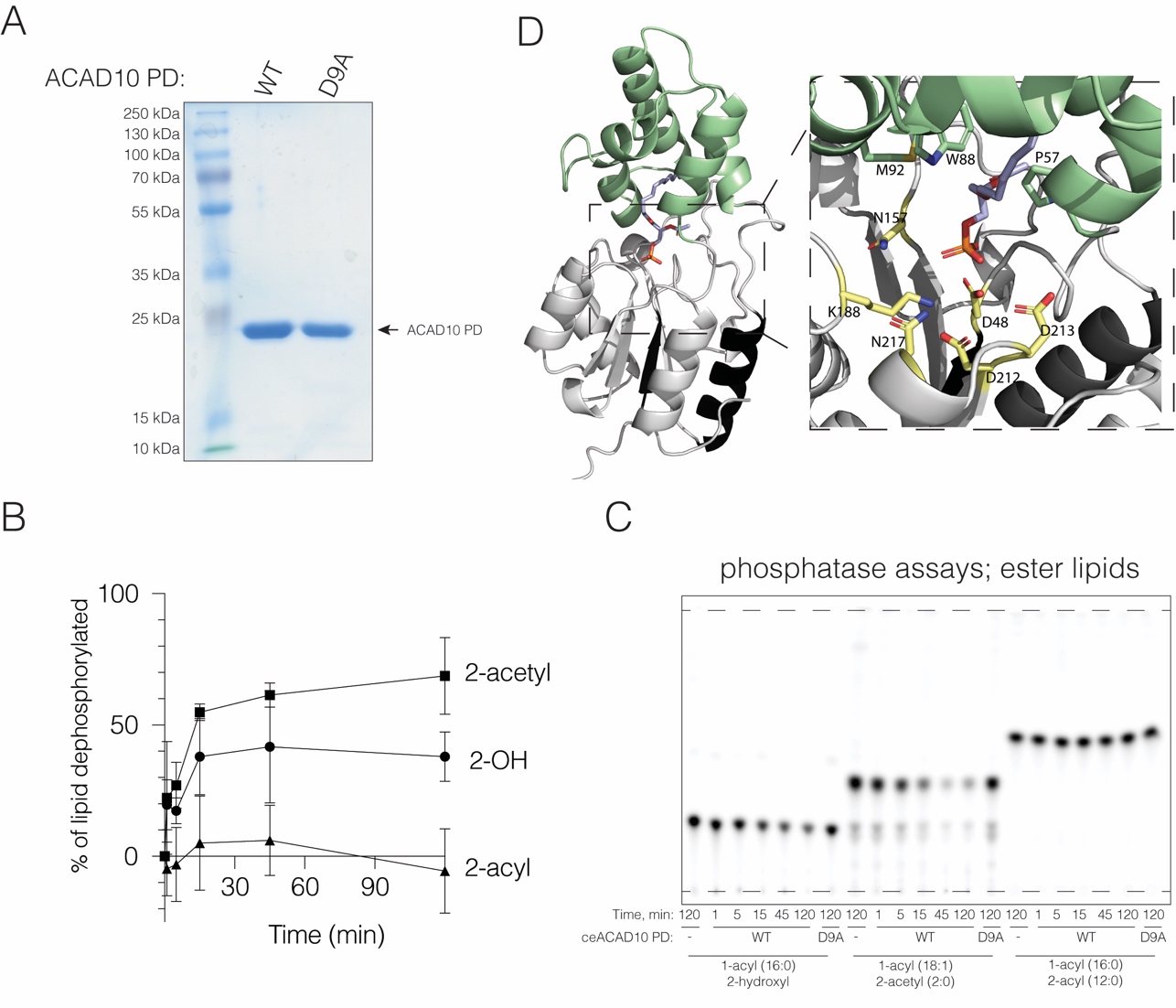

**Figure S3: The C. elegans ACAD10 phosphatase domain dephosphorylates 1-alkylglycerol phosphate and 1-acylglycerol phosphate, related to Figure 3.**

**(A)** SDS-PAGE and Coomassie staining of recombinant *C. elegans* ACAD10 phosphatase domain.

**(B)** Quantification of Fig. 3C showing the time-dependent dephosphorylation of ^32^P-labelled 1-O-alkyl-2-hydroxyl-*sn*-glycero-3-phosphate (2-OH), 1-O-alkyl-2-acetyl-*sn*-glycero-3-phosphate (2-acetyl), and 1-O-alkyl-2-acyl-*sn*-glycero-3-phosphate (2-acyl) by *C. elegans* ACAD10 phosphatase domain Quantification of three independent experiments are shown. Data represent mean ± SD.

**(C)** Thin-layer chromatograms showing time-dependent dephosphorylation of ^32^P -labeled acyl lipid substrates by the *C. elegans* ACAD10 phosphatase domain (PD) or the inactive D9A mutant (DA). Reaction products were separated by TLC and visualized by phosphorimaging. Data are representative of at least three independent experiments.

**(D)** AlphaFold3 model of the *H. sapiens* ACAD10 phosphatase domain (PD; left) and a magnified view of the active site (right) depicting a docked 1-hexadecoxy-3-phosphonooxypropan-2-yl acetate (light blue). The five-helical C1 cap (green) is predicted to form part of the substrate-binding pocket and may contribute to substrate recognition. The predicted interaction between the phosphatase domain and lipid substrate is high confidence (chain pair ipTM=0.86).

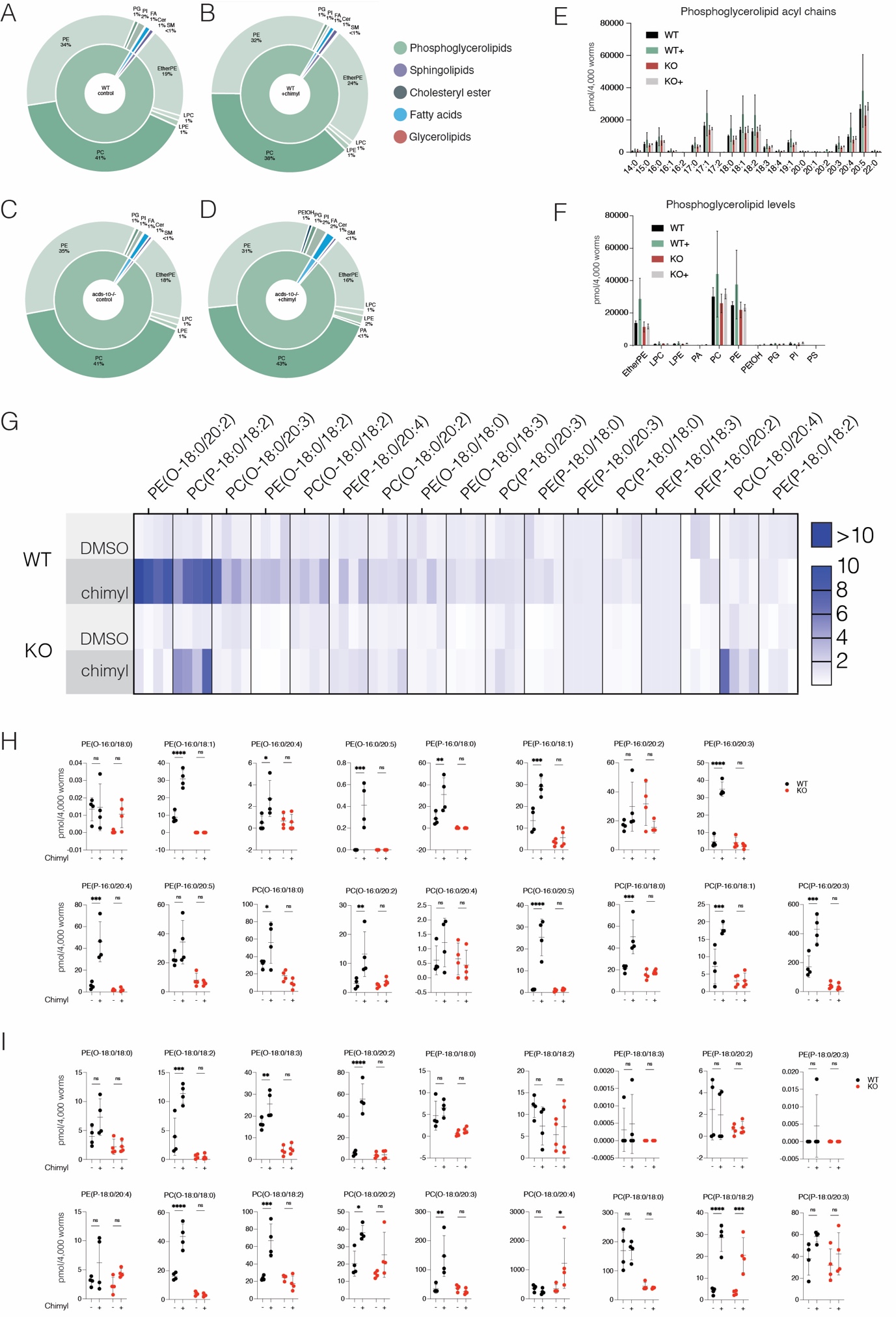

**Figure S4: Global lipidomic profiling in *C. elegans*, related to Figure 4.**

**(A-D)** Lipid class analysis of WT *C. elegans* **(A)**, WT *C. elegans* following dietary chimyl alcohol supplementation **(B)**, *acds-10-/-* *C. elegans* **(C)**, and *acds-10-/-* *C. elegans* following dietary chimyl alcohol supplementation **(D).** FA–Fatty Acids, Cer–Ceramides, SM–Sphingomyelins, LPC–Lysophosphatidylcholine, LPE–Lysophosphatidylethanolamine, PG–Phosphatidylglycerol, PI–Phosphatidylinositol, PA–Phosphatidic Acid.

**(E)** Plots showing the major detected phospholipid classes, presented as pmol per 4,000 worms, mean ± SD, n=4 (from 4,000-worms per experiment).

**(F)** Phospholipid acyl chain analysis, presented as pmol per 4,000 worms, mean ± SD, n=4 (from 4,000 worms per experiment).

**(G)** Heatmap depicting 18:0 alkyl/plasmenyl chain ether lipid levels in WT and *acds-10-/-* *C. elegans* under normal dietary conditions and after chimyl alcohol supplementation. Each cell represents the fold change for an individual replicate of 4,000 worms, normalized to the average value of the unfed WT condition.

**(H-I)** Individual ether lipids with (O/P-16:0) (**H**) or (O/P-18:0) alkyl/plasmenyl chains (**I**) in WT and *acds-10-/-* *C. elegans* after dietary chimyl alcohol supplementation (+). Data represent mean ± SD, points represent biological replicates of 4,000 worms per group, n=4. Data was analyzed using two-way ANOVA with Šídák multiple comparisons test ns: p>0.05, *p<0.05, ***p<0.001, ****p<0.0001.

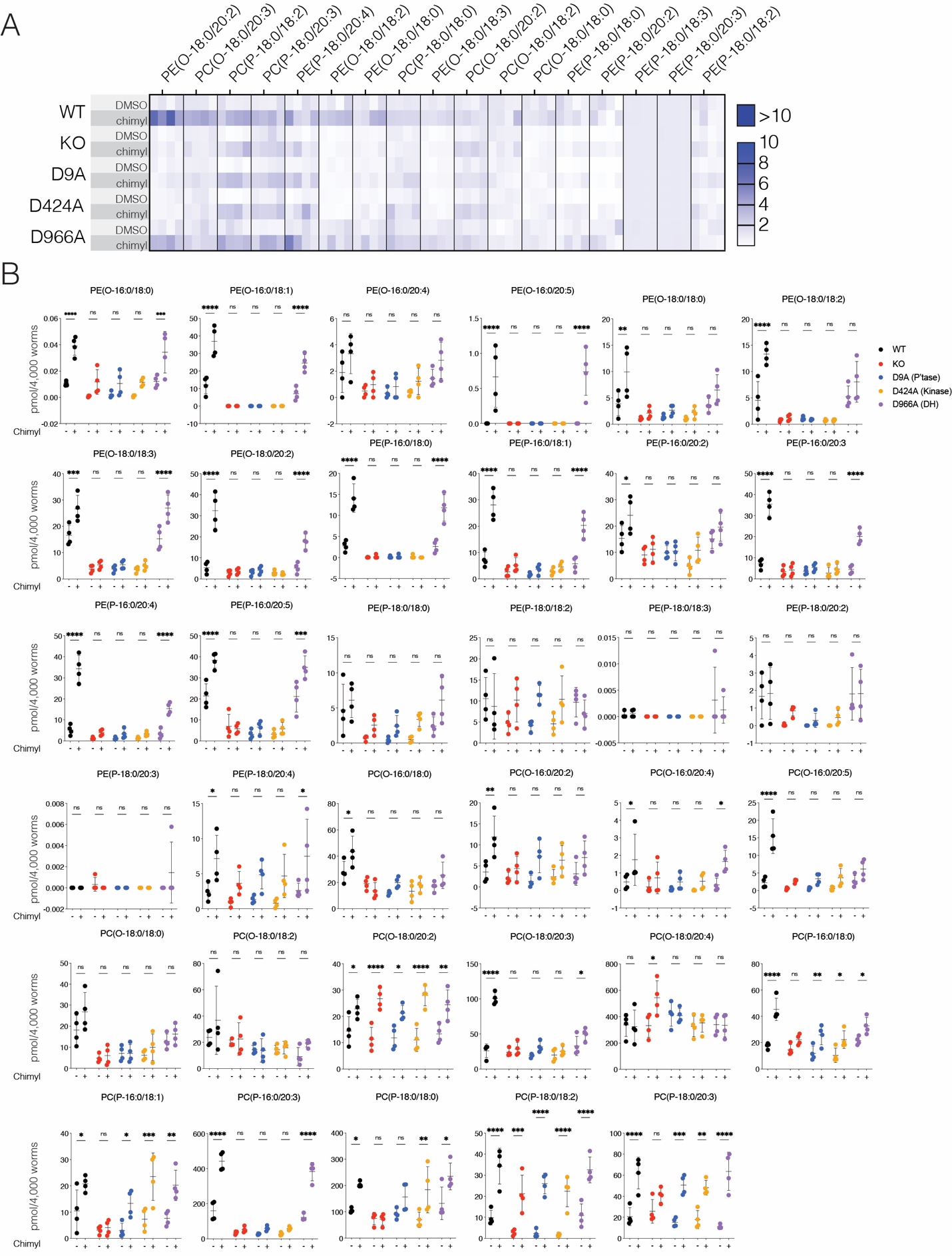

**Figure S5. Analysis of ether lipid levels in *C. elegans* strains supplemented with dietary chimyl alcohol, related to Figure 5**

**(A)** Heatmap depicting the levels of 18:0 alkyl/plasmenyl chain ether lipids in WT, *acds-10-/-*, and *C. elegans* strains harboring mutations in the phosphatase (D9A), kinase (D424A), and dehydrogenase (D966A) domains under normal dietary conditions and after dietary supplementation with chimyl alcohol. Each cell represents the fold change for an individual replicate of 4,000 worms, normalized to the average value of the unfed WT condition.

(**B**) Individual ether lipids with (O/P-16:0) and (O/P-18:0) alkyl/plasmenyl chains in WT, *acds-10-/-* and *C. elegans* strains harboring mutations in the phosphatase (D9A), kinase (D424A), and dehydrogenase (D966A) domains under normal dietary conditions and after dietary supplementation with chimyl alcohol (+). Data represent mean ± SD, points represent biological replicates of 4,000 worms per group, n=4. Data was analyzed using two-way ANOVA with Šídák multiple comparisons test ns; p>0.05, *p<0.05, **p<0.01, ***p<0.001, ****p<0.0001.

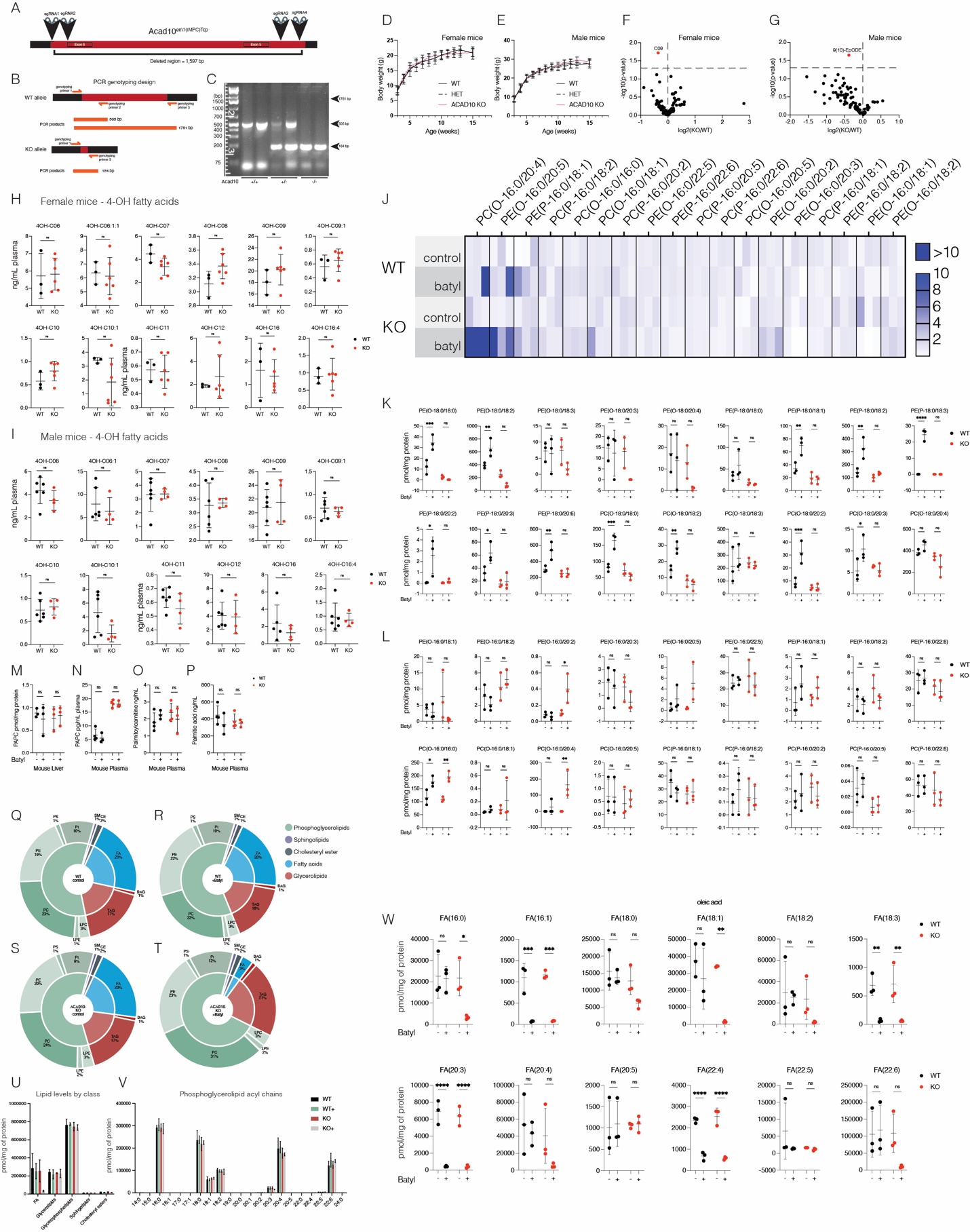

**Figure S6: Global lipidomic profiling in Acad10 KO mice, related to Figure 6**

**(A)** Diagram showing the mouse *Acad10* genomic locus targeted by sgRNAs used to make the Acad10 KO (*Acad10*^em1(IMPC)Tcp^) from International Mouse Phenotyping Consortium. The deleted region is marked in red.

**(B)** Predicted PCR products for Acad10 KO mouse genotyping. Expected PCR product sizes from WT and *Acad10* KO are depicted in orange.

**(C)** Agarose gel showing the PCR products for genotyping.

**(D-E**) Acad10 KO, WT, and heterozygous (HET) mice growth chart for female **(D)** and male **(E)** mice. Female mice: WT n=10, HET n=19, Acad10 KO n=19, male mice: WT n=8, HET=20, Acad10-KO n=11.

**(F-G)** Volcano plots showing hydroxylipids of female **(F)** and male **(G)** mice. Points depict average value of biological replicates, for female mice: WT n=3, ACAD10 KO n=6, male mice: WT n=6, ACAD10 KO n=4. P-values were generated using two-tailed unpaired t-tests.

**(H-I)** Individual 4-hydroxy fatty acid species in female **(H)** and male **(I)** mice. Graphs marked # had samples with fatty acid concentrations below lipidomics detection limit and were assigned 0 ng/mL. Data represent mean ± SD, points represent individual mice, for female mice: WT n=3, Acad10 KO n=6, male mice: WT n=6, KO=4. Data was analyzed using a two-tailed unpaired t-test, ns: p>0.05, **p<0.01.

**(J)** Heatmap showing 16:0 alkyl/plasmenyl chain ether lipid levels in WT and Acad10 KO mice under normal dietary conditions and after batyl alcohol supplementation. Each cell represents the fold change for an individual mouse, normalized to the average value of the unfed WT condition

**(K-L)** Individual ether lipids with (O/P-18:0) (**K**) or (O/P-16:0) alkyl/plasmenyl chains (**L**) in WT and Acad10 KO mice after dietary batyl alcohol supplementation (+)**.** Data represent mean ± SD, points represent individual mice, n = 3. Data was analyzed using two-way ANOVA with Šídák multiple comparisons test, ns: p>0.05, *p<0.05, ***p<0.001, ****p<0.0001.

**(M-N)** Liver **(M)** and plasma **(N)** 1-palmitoyl-2-acetyl-*sn*-glycero-3-phosphatidylcholine (PAPC) levels in WT and Acad10 KO mice after dietary batyl alcohol supplementation (+). Data represent mean ± SD, points represent individual mice, n=3 mice per group. Data was analyzed using two-way ANOVA with Šídák multiple comparisons test, ns: p>0.05.

**(O-P)** Plasma palmitoylcarnitine **(O)**, and palmitic acid **(P)** levels in WT and Acad10 KO mice after dietary batyl alcohol supplementation (+). Data represent mean ± SD, points represent individual mice, n=3. Data was analyzed using two-way ANOVA with Šídák multiple comparisons test, ns: p>0.05.

**(Q-T)** Lipid class analysis of liver samples from WT mice **(Q)**, WT mice supplemented with dietary batyl alcohol **(R)**, Acad10 KO mice **(S)**, and Acad10 KO mice supplemented with dietary batyl alcohol **(T)**. FA–Fatty Acids, Cer–Ceramides, SM–Sphingomyelins, LPC–Lysophosphatidylcholine, LPE–Lysophosphatidylethanolamine, PG–Phosphatidylglycerol, PI–Phosphatidylinositol, PA–Phosphatidic Acid, PS–phosphatidylserine.

**(U)** Individual major liver lipid class comparisons between genotypes and treatments. Data represent mean ± SD, n=3.

**(V)** Individual liver phospholipid acyl chain analysis between genotypes and treatments. Data represent mean ± SD, n=3.

**(W)** Liver free fatty acid levels in WT and Acad10 KO mice following dietary batyl alcohol supplementation. Data represent mean ± SD, points represent individual mice, n=3. Data was analyzed using two-way ANOVA with Šídák multiple comparisons test, ns: p>0.05, *p<0.05, **p<0.01, ***p<0.001, ****p<0.0001.

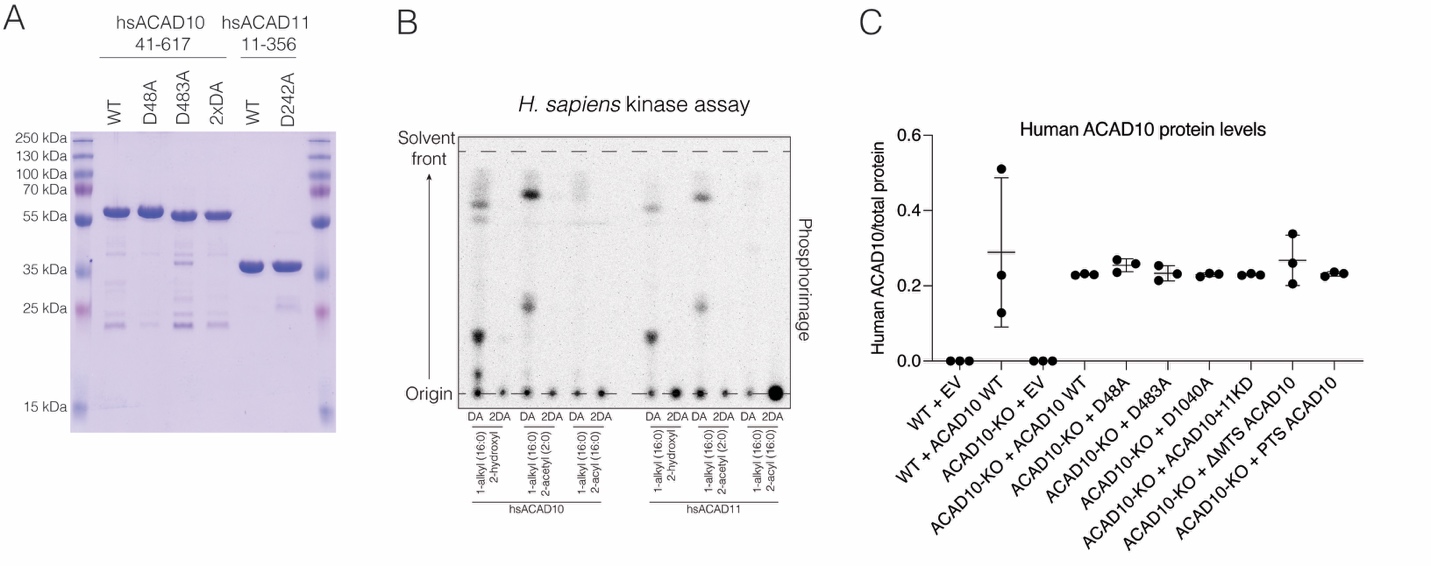

**Figure S7: Human ACAD10 and ACAD11 function in ether lipid metabolism, related to Figure 7.**

**(A)** SDS-PAGE and Coomassie staining of recombinant human ACAD10 phosphatase and kinase domain fragment (41-617) and ACAD11 kinase domain fragment (11-356).

**(B)** Thin-layer chromatogram showing incorporation of ^32^P from [ γ-^32^P]ATP into chimyl alcohol or ether lipids containing either an acetyl or long-chain acyl group at the sn-2 position by *H. sapiens* ACAD10 (hsACAD10; residues 41–617), which contains both the phosphatase domain (PD) and kinase domain (KD) and the ACAD11 kinase domain (hsACAD11). Reaction products were separated by TLC and visualized by phosphorimaging. DA, D48A; 2DA, D48A/D483A.

**(C)** Proteomic analysis of human ACAD10 expression in WT and Acad10-KO Hepa1-6 cells expressing the indicated human ACAD10 constructs. Human ACAD10 peptide abundance was quantified by LC-MS/MS and normalized to total protein concentration. Endogenous mouse ACAD10 was not detected, as indicated by the empty vector (EV) condition. Data represent mean ± SD.

**Table S1. Data collection and refinement statistics, ACAD10:Mg^2+^-AMP-PNP structure.**

| **Data collection** | |
| --- | --- |
| Space group | P2_1_ |
| Cell constants (Å) | a = 94.97, b = 95.89, c = 95.54 β = 114.09° |
| Wavelength (Å) | 0.97946 |
| Resolution range (Å) | 36.89 – 2.15 (2.19 – 2.15) |
| Unique reflections | 84,386 (4,191) |
| Multiplicity | 4.5 (3.5) |
| Data completeness (%) | 99.5 (98.8) |
| *R*_merge_ (%)^b^ | 10.5 (104.3) |
| *R*_pim_ (%)^c^ | 4.8 (52.4) |
| I/σ(I) | 15.8 (1.5) |
| CC_1/2_ (highest resolution shell) | 0.49 |
| Wilson B-value (Å^2^) | 25.8 |
| **Refinement statistics** | |
| Resolution range (Å) | 36.89 – 2.15 (2.20 – 2.15) |
| No. of reflections *R*_work_/R_free_ | 79,215/2,002 (3,112/82) |
| Data completeness (%) | 93.2 (53.0) |
| Atoms (non-H protein/nucleotide/water) | 21,542/124/413 |
| *R*_work_ (%) | 18.5 (23.9) |
| *R*_free_ (%) | 22.2 (25.8) |
| R.m.s.d. bond length (Å) | 0.003 |
| R.m.s.d. bond angle (°) | 0.59 |
| Mean B-value (Å^2^) (chain A/chain B/chain C/chain D/ligands/waters) | 34.2/47.4/38.4/51.8/35.8/33.3 |
| Ramachandran plot (%) (favored/additional/disallowed)^d^ | 96.2/3.65/0.15 |
| Clashscore^d^ | 2.5 |
| Maximum likelihood coordinate error | 0.21 |
| Missing residues | A: 1-2, 325-330, 353-355. B: 1-3, 67-71, 266-268, 323-334, 352-355. C: 1-3, 69-71, 188-189, 266-269, 326-330. D: 1-5, 67-71, 267, 325-333, 354-355. |

Data for the outermost shell are given in parentheses.

^a^Bijvoet-pairs were kept separate for data processing.

^b^*R*_merge_ = 100 Σ_h_Σ_i_|*I_h,i_*— 〈*I_h_*〉*|/*Σ*_h_*Σ_i_ 〈*I_h,i_*〉, where the outer sum (h) is over the unique reflections and the inner sum (i) is over the set of independent observations of each unique reflection.

^c^ *R*_pim_ = 100 Σ_h_Σ_i_ [1/(n_h_ - 1)]^1/2^|*I_h,i_*— 〈*I_h_*〉*|/*Σ*_h_*Σ_i_ 〈*I_h,i_*〉, where n_h_ is the number of observations of reflections **h**.

^d^As defined by the validation suite MolProbity ^49^.

STAR Methods

1. Experimental model and study participant details

**Bacterial strains**

For protein purification BL21 (DE3) or Rosetta (DE3) *E. coli* were used as indicated. Bacteria were grown in LB or Terrific broth (TB) as indicated at 37^o^C with rocking.

For *C. elegans* maintenance OP50 and HT115 *E. coli* strains were used as indicated.

**Hepa1-6 cell generation**

For normal propagation, Hepa1-6 (mouse hepatoma; ATCC; CRL-1830) cells were cultured in high-glucose DMEM containing 10% fetal bovine serum (FBS) (Gibco; Lot # 2098782) at 37°C and 5% CO_2_. Monoclonal CRISPR knockout Hepa1-6 cell lines were generated by the GEiC at Washington University using two different single-stranded guide RNAs that targeted early exons of *Acad10* and *Acad11*. After selection and verification by next-generation sequencing, two monoclonal lines per gene knockout combination were chosen for experimentation. Hepa1-6 cells used in our experiments were passaged no more than 10 times. Cell lines were verified to be mycoplasma free through use of the LookOut Mycoplasma qPCR detection kit (Sigma-Aldrich).

***C. elegans* strains and details**

Nematodes were maintained at 15°C on standard nematode growth medium (NGM) agar plates seeded with *E. coli* OP50. All experimentation was performed using hermaphrodites at 20°C. SUNY Biotech generated via CRISPR Cas-9 the four *acds-10* mutant strains: the null, phosphatase (D9A), kinase (D424A), and dehydrogenase (D966A) domain mutants. Mutant strain protein levels were validated via western blot using α-acds-10 antibodies generated in this study. The N2 strain from the *Caenorhabditis* Genetics Center (CGC) was utilized as the control in all experimentation.

**Mouse generation and husbandry**

All animal work was approved by the Institutional Animal Care and Use Committee of the College of Agricultural and Life Sciences at the University of Wisconsin-Madison. Mice were housed with 12-hour light-dark cycles, at 35% humidity 21°C. Mice had ready access to food and water, unless fasting. Cryopreserved *Acad10*^-/-^ mouse sperm was purchased from the Canadian Mutant Mouse Repository (*Acad10*^em1(IMPC)Tcp^). *In vitro* fertilization of C57BL/B6J donor eggs with the sperm was performed by the UW-Madison Animal Models Core. Two-cell embryos were implanted in C57BL/B6J recipient mothers, and the offspring were genotyped to identify *Acad10^-/+^*. Breeder mice were maintained on a high-energy diet (Formulab Diet 5015). Experimental mice were maintained on standard chow diet (Formulab Diet 5008). For oxylipin analysis, at approximately 4-month-old mice were fasted for 6 hours, euthanized and had plasma and tissues harvested (Male WT n=6; Male KO n=4; Female WT n=3; Female KO n=6). For ether lipid and PAF quantification, mice were fed either standard chow (Envigo; TD.00606) or batyl alcohol-supplemented chow (customized Envigo; TD.00606 with addition of 2% w/v batyl alcohol^50^.

**Human Demographics and Sample Collection**

All human studies were overseen by the University of Wisconsin Madison Institutional Review Board (#2023-0230). Biological samples from Akimel O'odham Population were accessed through the Data for Indigenous Implementations, Interventions, and Innovations Tribal Data Repository (D4I TDR), a federally funded, Indigenous-led repository. Data access and use were governed under the CARE Principles for Indigenous Data Governance (Collective Benefit, Authority to Control, Responsibility, and Ethics), which the repository's operating framework centers alongside standard FAIR data practices to recognize and empower Indigenous Peoples' rights and interests in their own data. Consistent with this framework, participating Tribes directed the terms under which their data could be accessed and used, and sample access for this analysis was reviewed and approved in accordance with the D4I TDR Data Access and Use Committee's governance procedures and the Tribal Nation's own data-sharing terms.

All samples were obtained from the repository in de-identified form. Plasma was processed from blood collected with sodium heparin tubes (BD367874). Tubes were inverted 5-10 times immediately after collection, and centrifuged at 1200 x g for 10 minutes at 4°C. Plasma was dispensed in 1mL aliquots for storage at -80 °C. Demographic variables (e.g., age, sex, household or geographic detail) were blind matched by the D4I TDR with age and sex matching. Genetic polymorphisms of rs601663 and rs659964 are preserved within the D4I TDR for access after Tribal approval. Because the Akimel O'odham community constitutes a small, geographically and socially bounded population, even coarse demographic descriptors can substantially increase the risk of re-identifying individuals or family groups when cross-referenced with other public or semi-public records. Demographic variables were therefore withheld from the dataset as a condition of data governance approval. This approach follows the Tribal data-governance principle that data stewardship and disclosure risk are determined by the originating Tribal Nation, per D4I TDR policy.

2. **Method details**

**Generation of plasmids**

*C. elegans* ACAD10 (acds-10) phosphatase domain (amino acids 1-212) or kinase domain (amino acids 209-568) were PCR amplified from *C. elegans* cDNA and cloned into ppSumo (a modified pET-28a based vector containing a 6XHis-SUMO tag and the yeast smt3 CDS). *E. coli* codon optimized *Caldalkalibacillus thermarum* ACAD10 (WP_007504381.1), and *Myxococcus xanthus* ACAD10 (Q1CZY9_MYXXD) were synthesized as gBlocks (IDT), while *E. coli* codon optimized *Marinicauda pacifica* (A0A4S2HBA5-9PROT), *Caldilinea aerophila* strain DSM 14535 (I0I2H8_CALAS), *Aeromicrobium* sp. IC_218 (A0A4V2NSZ3_9ACTN), *Aggregicoccus* sp. 17bor-14 (A0A6I2GPB5_9BACT), and *Serpentinimonas raichei* (A0A060NI31_9BURK) ACAD10s were synthesized by Twist Bioscience and cloned into ppSumo vector.

*E. coli* DgkA was PCR amplified from DH5α *E. coli* gDNA and cloned into the ppSumo vector.

All mutations were made using QuikChange site directed mutagenesis, and primers were designed with the online Agilent QuikChange Primer Design website.

**Protein expression and purification**

*C. elegans* ACAD10 kinase domain (209-568) WT and D424A constructs in ppSUMO plasmid were transformed into Rosetta (DE3) *E. coli*. Cells were grown in 1 L LB in the presence of 50 μg/ml kanamycin at 37^o^C and protein expression was induced at an OD_600_ of ~0.8 with 0.4 mM IPTG. Proteins were expressed overnight at 18^o^C. Cultures were centrifuged at 3,000 x g for 10 minutes, and the bacterial pellets were resuspended in lysis buffer (50 mM Tris-HCl pH 8.0, 300 mM NaCl, 1 mM DTT, 25 mM imidazole, 1 mM PMSF). Resuspended cells were lysed by sonication at 70% amplitude for 2:30 min with 5 sec ON/ 15 sec OFF pulses and the lysates were cleared by centrifugation at 30,000 x g for 30 minutes. The lysate was incubated with HisPur™ Ni-NTA resin (0.5 mL of packed resin per 1L prep, Thermo Fisher Scientific 88222), pre-washed with lysis buffer. The resin was washed in 50 mM Tris-HCl pH 8, 300 mM NaCl, 25 mM imidazole, 1 mM DTT and the proteins were eluted with elution buffer 50 mM Tris-HCl, pH 8.0, 300 mM NaCl, 300 mM imidazole, 1 mM DTT and cleaved overnight at 4^o^C with the Sumo protease, Ulp. The cleaved proteins were further purified by size-exclusion chromatography using a Superdex HiLoad S75 16/600 (Cytiva) or Superdex S75 increase 10/300 (Cytiva) size exclusion columns into size exclusion buffer (50 mM Tris, pH 8.0, 300 mM NaCl, 1 mM DTT). ACAD10 was concentrated to 1 mg/mL, aliquoted, flash frozen and stored at -80 ^o^C until use.

Bacterial ACAD10 kinase domain homologs *Marinicauda Pacifica* (A0A4S2HBA5-9PROT), *Caldilinea aerophila* strain DSM 14535 (I0I2H8_CALAS), *Aeromicrobium* sp. IC_218 (A0A4V2NSZ3_9ACTN), *Aggregicoccus* sp. 17bor-14 (A0A6I2GPB5_9BACT), *Myxococcus xanthus* (Q1CZY9_MYXXD), and *Serpentinimonas raichei* (A0A060NI31_9BURK) were purified using the same steps as *C. elegans* ACAD10 kinase domain.

All *C. thermarum* ACAD10 kinase purification steps were carried out at room temperature including overnight protein expression due to its origin from a thermophile.

*E. coli* DgkA kinase purification followed the same steps as *C. elegans* ACAD10 kinase domain, but the Ulp cleavage step was skipped and the final enzyme used in assays contained the 6His-SUMO tag.

*C. elegans* ACAD10 phosphatase domain (1-212) WT and D9A proteins were purified following the same steps as described above but all the buffers included 1 mM MgCl_2_. After size-exclusion chromatography the protein was diluted to reduce the NaCl concentration to 50 mM or lower and it was then run on an anion-exchange column (HiTrap™ Capto™ Q 5 mL) using a linear gradient from 50 mM Tris-HCl pH 8, 25 mM NaCl, 1 mM MgCl_2_, 1 mM DTT to 1 mM Tris-HCl pH 8, 500 mM NaCl, 1 mM MgCl_2_, 1 mM DTT to remove any remaining contaminating protein.

*Human ACAD10 (41-617, E. coli* codon optimized*) and ACAD11 (11-356)* WT and mutants were purified using the same steps as above but BL21 (DE3) *E. coli* were used and grown in Terrific broth (TB), and all the buffers included 1 mM MgCl_2_. After incubation with SUMO protease Ulp, the protein was diluted to reduce the NaCl and imidazole concentration to 50 mM or lower and it was then run on an anion-exchange column (HiTrap™ Capto™ Q 5 mL) using a linear gradient from 50 mM Tris-HCl pH 8, 25 mM NaCl, 1 mM MgCl_2_, 1 mM DTT to 1 mM Tris-HCl pH 8, 500 mM NaCl, 1 mM MgCl_2_, 1 mM DTT. Peaks containing ACAD10/11 were further purified by size-exclusion chromatography using Superdex S75 increase 10/300 (Cytiva) size exclusion columns into size exclusion buffer (50 mM Tris, pH 8.0, 300 mM NaCl, 1 mM DTT). Protein was concentrated to 1 mg/mL, aliquoted, flash frozen and stored at -80 ^o^C until use.

**Preparation of crude mitochondrial extract by differential centrifugation**

The method was adapted from previous isolation protocols for crude mitochondrial extraction ^51,52^. Two fresh mouse livers were placed in a 50 mL beaker with enough MSHE+BSA buffer (210 mM mannitol, 70 mM sucrose, 5 mM HEPES, 1 mM EGTA, pH 7.2 with KOH supplemented with 0.2% w/v fatty acid-free (Fraction V) BSA) to cover the tissue and the whole procedure was carried out on ice. The tissue was minced with scissors and the MSHE+BSA buffer was changed a few times until it was clear of blood. The tissue was then homogenized using the Fisherbrand™ 150 Homogenizer (Fisher Scientific, 15-340-168) on ice until the homogenate was clear of larger debris. More MSHE+BSA was added to reach 40 mL volume in the 50 mL tube. The samples were then centrifuged at 12,000 g for 10 min at 4^o^C. The top layer of floating lipids was removed, and the supernatant was gently aspirated. The pellet was resuspended with MSHE+BSA buffer and spun again at 800 x g for 5 min at 4^o^C to pellet nuclei and tissue debris. The supernatant was then centrifuged at 12,000 x g for 10 min at 4^o^C. The pellet, that contains mitochondria as well as plasma membrane, endoplasmic reticulum, and mitochondria-adjacent membranes ^51^, was resuspended in 500 µL of MSHE+BSA. The sample was gently mixed by pipetting, and an additional 500 µL of MSHE+BSA was added, followed by centrifugation at 12,000 x g for 10 min at 4^o^C. The supernatant was aspirated, and the pellet was resuspended in 500 µL of MSHE without the BSA. The sample was gently mixed, then centrifuged at 12,000 x g for 10 min at 4^o^C. The pellet was resuspended in ~100 µL of MSHE without BSA. Protein concentrations were measured using Bradford’s reagent. Membranes were adjusted to 5 mg/mL, aliquoted to avoid multiple freeze-thaws, flash-frozen using liquid nitrogen, and stored at -80^o^C until use.

**Kinase activity assays using crude mitochondrial extract**

For *C. elegans* ACAD10 kinase domain, kinase assays were carried out with 2 µg of crude mitochondrial fraction (by protein concentration) in 50 mM Tris pH 8, 1 mM DTT, 5 mM MgCl_2_, 1 mM acetyl-CoA, 1 mM [γ-^32^P]ATP (specific activity = 500 cpm/pmol), 40 mM NaF unless specified otherwise. For coenzyme A derivative screen, each CoA derivative was used at 0.1 mM concentration. Reactions were carried out at 37 ^o^C for 15 min unless specified otherwise. All assays were carried out in 20 µL volume, reactions were started with the addition of the kinase protein and stopped with 500 µL of MeOH.

For the SDS-PAGE experiment, reactions were incubated for 2h at 37^o^C. Reaction was stopped with 5x SDS-PAGE loading dye, 1:5 ratio. 1:100 diluted Proteinase K (stock ~20mg/mL, Thermo Scientific, EO0491) was added 1 µL per reaction and was incubated for 1 h at 37^o^C to digest all protein.

**Kinase activity assays using 1-O-alkylglycerols**

Lipids were resuspended in 100% ethanol to a concentration of 2 mM. For *C. elegans* ACAD10 kinase domain, kinase assays were carried out with 0.1 mM of substrate lipids in 50 mM Tris pH 8, 1 mM DTT, 5 mM MgCl_2_, 1 mM acetyl-CoA, 0.5 mM [γ-^32^P]ATP (specific activity = 2000 cpm/pmol) for 2 h at 37 ^o^C unless specified otherwise.

For ACAD10 kinase domain homolog assays, conditions were as above with a few exceptions. 1 mM [γ-^32^P]ATP was used (specific activity = 500 cpm/pmol). *Marinicauda Pacifica* (A0A4S2HBA5-9PROT), *Caldilinea aerophila* strain DSM 14535 (I0I2H8_CALAS), *Aeromicrobium* sp. IC_218 (A0A4V2NSZ3_9ACTN), *Aggregicoccus* sp. 17bor-14 (A0A6I2GPB5_9BACT) were incubated for 1 h at pH 8 at 37 ^o^C. *Myxococcus xanthus* (Q1CZY9_MYXXD) was incubated for 30 min at pH 8 at 37 ^o^C. *Caldalkalibacillus thermarum* (WP_007504381.1) was incubated for 15 min at pH 8 at 60 ^o^C. *Serpentinimonas raichei* (A0A060NI31_9BURK) was incubated for 1 h at pH 10 at 30 ^o^C.

All assays were carried out in 20 µL reaction volumes, and reactions were started by the addition of the kinase and stopped with 500 µL of MeOH.

**Saponification of lipids**

Kinase reactions for saponification experiments were stopped with 50 µL of MeOH. For samples designated for saponification 500 µL of 0.5 M KOH (dissolved in MeOH) was added; for the remaining samples, 500 µL of MeOH was added instead. All samples were heated at 80^o^C for 45 min. Samples were allowed to cool to room temperature and the lipid extraction was performed.

**Extraction of lipids**

Lipid extractions were performed in accordance with the original protocol ^53^. The kinase or phosphatase reactions were terminated by the addition of 500 µL of MeOH. Then, 500 µL of chloroform, 250 µL of water, and 5 µL of 36.5-38% HCl (VWR BDH3026-500MLP) were added to induce phase separation. For saponified samples, an extra 20 µL of 36.5-38% HCl was added to neutralize the KOH and achieve acidification of the sample. Addition of HCl protonates phosphorylated lipids to increase solubility in the nonpolar organic phase. The mixture was vortexed and then centrifuged at 3500 x g at room temperature for 5 minutes in a table-top centrifuge to give a two-phase system (aqueous top, organic bottom). The bottom phase was recovered by inserting a pipette tip through the upper phase with gentle positive pressure to avoid contamination from the aqueous phase and then withdrawing the bottom phase, making sure to avoid the interface or the upper phase. To prepare the “authentic upper phase,” a 1:1:1 chloroform:methanol:water mixture was vortexed and centrifuged at 3500 x g at room temperature for 5 minutes. 500 µL of the authentic upper phase was transferred into the tube with the lipid extraction organic phase, vortexed, and separated by centrifugation at 3500 x g at room temperature for 5 minutes. For samples where [γ-^32^P]ATP concentration was higher (2000 cpm/pmol), an additional wash with the authentic upper phase was performed. The organic phase was once again transferred to a new tube and dried in a vacuum concentrator (Eppendorf 022820109) with V-AL mode, 30^o^C for 40 minutes.

**Preparation of ^32^P-labelled substrates for phosphatase assays**

Lipids for kinase assays were resuspended in 100% ethanol to a concentration of 10 mM. Lipids were diluted in reaction buffer to a final concentration of 0.5 mM and 5% ethanol. The final reaction buffer was made up of 50 mM Tris pH 8, 1 mM DTT, 5 mM MgCl_2_, 1 mM [γ-^32^P]ATP (specific activity = 500 cpm/pmol). Reactions were started by the addition of the lipid kinase. 1-O-hexadecyl-*rac*-glycerol, 1-O-hexadecyl-2-O-acetyl-*sn*-glycerol, 1-O-hexadecyl-2-palmitoyl-*rac*-glycerol, 1-hexadecyl glycerol, 1-oleoyl-2-acetyl-*sn*-glycerol, and 1-palmitoyl-2-lauroyl-*rac*-glycerol were phosphorylated by *E. coli* DgkA. The reaction was stopped by the addition of 500 µL MeOH and 500 µL of chloroform. To separate phosphorylated from non-phosphorylated lipids, 250 µL of 1 mM NaOH in ddH_2_O was added; the samples were vortexed and centrifuged for 5 min at 3,500 x g at room temperature. The majority of the phosphorylated lipids reside in the aqueous phase, while the non-phosphorylated lipids were retained in the organic phase (chloroform). The aqueous phase was transferred to a new tube with fresh 500 µL of chloroform, and 250 µL of 0.7% HCl in deionized H_2_O was added, acidifying the extract and protonating the phosphorylated lipids. Samples were vortexed and centrifuged for 5 min at 3,500 x g. The protonated phosphorylated lipids were now transferred into the organic phase. The organic phase was transferred into 500 µL of an authentic upper phase from chloroform:MeOH:deionized water mixture, vortexed, and centrifuged for 5 min at 3,500 x g. The authentic upper phase was prepared by vortexing a 1:1:1 ratio of chloroform:MeOH:deionized H_2_O, centrifuging for 5 min at 3,500 x g, and separating the aqueous (upper) phase. The organic phase was transferred to a new tube with 500 µL of authentic upper phase, vortexed, and centrifuged for 5 min at 3,500 x g. The organic phase was transferred again to a new tube and dried in a vacuum concentrator with V-AL mode, 30 °C for 40 minutes.

**Phosphatase activity assays using ^32^P-labelled substrates**

Unless specified otherwise, phosphatase assays were performed for 15 min at 37^o^C in 20 µL reactions containing 1 µL of ^32^P-labelled phosphorylated lipids resuspended in ethanol (200-300 cpm and 5% ethanol per reaction), 50 mM Tris pH 7.5, 1 mM MgCl_2._ The reactions were started by the addition of 0.5 µM of *C. elegans* ACAD10 phosphatase domain. For the substrate specificity assays, final concentrations of 0.5 µM POPC lipid were included in the reaction. Reactions were stopped by the addition of 500 µL of MeOH and proceeding with the lipid extraction without the second wash step with the authentic upper phase.

**Thin layer chromatography**

The dried organic phase samples were resuspended in 4.5 µL of TLC buffer (10:4:3:2:1 chloroform:acetone:methanol:acetic acid:water) and spotted onto Silica TLC plates (Silica gel 60 F₂₅₄, L × W 20 cm × 20 cm, glass support, Supelco 1.05715). Plates were left to dry then placed in a glass chamber filled, ~ 1 cm deep, with TLC buffer. Once the solvent front reached around 2/3rds the way to the top of the plate, the plate was removed from the chamber, the solvent front was marked in pencil, and the plate was left to dry. Once the TLC plate was fully dry, it was wrapped in a protective plastic sheet and placed in a cassette with a blanked Phosphor screen (Cytiva, 28956475). The phosphor screen was stored on the TLC plate in the dark overnight or for two nights and imaged with Typhoon biomolecular imager (Amersham).

**Crystallization, data collection and structure determination**

*C. thermarum* ACAD10 was prepared by expression and purification from *E. coli* as described above and concentrated to 10 mg/mL in 10 mM Tris-HCl pH 8.0, 150 mM NaCl and 1 mM TCEP. ACAD10:Mg^2+^-AMP-PNP was prepared by incubation of ACAD10 at 10 mg/ml with 5 mM MgCl_2_ and 1 mM AMP-PNP for 2 hours. ACAD10: Mg^2+^-AMP-PNP crystals were grown by the sitting drop vapor diffusion method at 20°C in 24-well Cryschem trays using a 1:1 ratio of protein/reservoir solution containing 11% w/v PEG 4,000, 0.2 M ammonium sulfate and 0.1 M sodium acetate pH 3.6. ACAD10:Mg^2+^-AMP-PNP crystals were cryo-protected with 13% (w/v) PEG 4,000, 0.2 M ammonium sulfate, 0.1 M sodium acetate pH 3.6, 5 mM MgCl_2_, 1 mM AMP-PNP and 30% (w/v) ethylene glycol, diffracted to a minimum Bragg spacing (dmin) of 2.15 Å and exhibited the symmetry of space group P2_1_ with cell dimensions of a=94.97 Å, b=95.89 Å, c=95.54 Å, β=114.09°, and contained four ACAD10:Mg^2+^-AMP-PNP per asymmetric unit. Diffraction data for ACAD10:Mg^2+^-AMP-PNP were collected at beamline 19-ID (SBC-CAT) at the Advanced Photon Source (Argonne National Laboratory, Argonne, Illinois, USA) and were processed in the program HKL-3000 ^54^ with applied corrections for effects resulting from absorption in a crystal and for radiation damage ^55,56^, the calculation of an optimal error model, and corrections to compensate the phasing signal for a radiation-induced increase of non-isomorphism within the crystal ^57,58^. Phases for ACAD10:Mg^2+^-AMP-PNP were calculated via molecular replacement within the program Phaser ^59^ using residues 1-355 of a model generated in AlphaFold2 as a search model ^60^. Editing of the model was performed by multiple cycles of manual rebuilding in the program Coot ^61^ and refinement in the program Phenix ^62^. Positional and isotropic atomic displacement parameter (ADP) as well as TLS ADP refinement was performed in the program Phenix with a random 2.53% of all data set aside for an R_free_ calculation. Data collection and structure refinement statistics are summarized in **Table S1.**

**Quantification of TLC signal for phosphatase assays**

^32^P-labelled lipids were quantified using ImageJ software. Only non-saturated signal was quantified. Phosphatase activity was measured as % phosphorylated lipid remaining compared to the no phosphatase control (100%). Results are presented as the mean +/- standard deviation of at least 3 independent replicates.

**Quantification of *C. elegans* fecundity**

Late-larval stage (L4) animals were manually transferred to individual 35 mm Nematode Growth Medium (NGM) plates seeded with *E. coli* OP50. Every 24 hours, for five consecutive days, the adult animals were transferred to new plates. Progeny were counted one day after the P0 animal was moved to a new plate to ensure sufficient time for eggs to hatch. Once hatched, progeny were counted by quadrant, along with any unhatched eggs. This experiment was conducted at 20°C.

***C. elegans* lifespan determination**

Lifespan analyses were performed on NGM/Carb plates at 20°C on HT115 *E. coli* expressing OP50. Animals were age-synchronized using hypochlorite treatment and plated on bacteria and grown from hatch to adulthood at 20°C. Ten adult worms were moved away from progeny onto ten fresh plates empty vector plates at Day 1, for a starting total of 100 adult worms per condition, and treated with 100 mg/ml 5-Floro-2'-deoxyuridine (FUDR) to prevent progeny development of eggs laid post-day 1 adulthood.

***C. elegans* lysate preparation**

Age synchronized populations of worms were cultivated on nematode growth media (NGM) agarose plates supplemented with HT115 E. coli at 20°C. Day 1 adult animals were washed off the plates with M9 buffer, centrifuged at 1000 x g for 30 s at room temperature and washed twice with M9 before being transferred to 1.5 ml Eppendorf tubes and rapidly flash frozen in liquid nitrogen.

Worm extracts were generated by glass bead disruption in non-denaturing lysis buffer [50 mM Hepes pH 7.4, 150 mM NaCl, 1mM EDTA, 1% Triton, EDTA-free mini-protease inhibitor cocktail (Roche), phosSTOP phosphatase inhibitor cocktail (Roche)]. Crude lysates were subjected to centrifugation at 7500 x g at 4°C for 5 min.

**Purification of rabbit polyclonal α-acds-10 antibodies**

*C. elegans* ACAD10 kinase domain was used to inoculate rabbits for generation of rabbit α-acds-10 anti-serum (Cocalico Biologicals). Total IgG was partially purified by ammonium sulfate precipitation ^63^. α-acds-10 antibodies were affinity-purified by coupling *C. elegans* ACAD10 kinase domain to a HiTrap NHS-activated HP column (Cytva, 17071601) essentially as described ^64^. Briefly, the column was washed with 6 mL of 1 mM HCl to remove storage buffer. *C. elegans* ACAD10 kinase domain was purified as described above, and buffer exchanged into 0.2 M NaHCO_3_ pH 8.3, 0.5 M NaCl. Three mL of ~3.5 mg/mL of protein was loaded back and forth onto the column around 1 mL/min for 30 min at RT. The column was washed with 1 mL of 10 mM Tris pH 7.5. To deactivate the column, it was washed with a 6 mL wash of buffer A (0.5 M ethanolamine, 0.5 M NaCl, pH 8.3) and 6 mL of buffer B (0.1 M acetic acid, 0.5 M NaCl, pH 4). A second wash with buffer A was performed and buffer A was left to incubate on the column for 30 min at RT before proceeding with a second buffer B wash. A third round of buffer A and buffer B washes was performed, followed by a wash with 10 mM Tris pH 7.5. The column was further washed with 10 mL of 100 mM glycine pH 2.5, and with 10 mL of 0.2 M sodium borate, 160 mM NaCl, pH 8.0.

15 mL of combined serum from 2 rabbits were precleared by a 10,000 x g spin for 30 min. The supernatant was transferred into a glass beaker on a mixer at 4 ^o^C. 35 mL of saturated ammonium sulphate was slowly added into the supernatant and was left to mix for 2 h to precipitate the protein. The precipitated serum was centrifuged at 10,000 x g for 30 min at 4 ^o^C. The supernatant was then poured off and the remaining protein were resolubilized in 15 mL of 0.2 M sodium borate, 160 mM NaCl, pH 8.0. The solubilized protein was slowly loaded onto the column. The column was washed with 20 mL of 0.2 M sodium borate, 160 mM NaCl, pH 8.0, 20 mL of 10 mM Tris pH 7.5, 0.5 M NaCl.

The bound antibody was eluted into a 15 mL tube containing 1 mL of 1M Tris pH 8.0 using 9 mL of 100 mM glycine pH 2.5. The neutral pH was verified using pH strips. The antibody was concentrated and buffer exchanged to PBS in an Amicon® Ultra Centrifugal Filter, 30 kDa MWCO (Millipore, UFC9030). Glycerol was added to a final concentration of 50%, and NaN_3_ to 0.1% for storage at -20 ^o^C. The yield was around 1 mg of purified antibodies.

**Immunoblotting**

Samples were separated by SDS-PAGE then transferred to nitrocellulose membranes in 48 mM Tris base, 39 mM glycine, and 20% ethanol for 90 min at a constant 90 V. Membranes were blocked using 5 % milk in TBS-T (20 mM Tris pH 7.5, 150 mM NaCl, 0.1% Tween-20) at RT for 1 h. Membranes were left incubating with the antibody diluted in 5% BSA in TBS-T overnight at 4 ^o^C. The next day, the membranes were washed 3 x 15 min with TBS-T at RT, and incubated with secondary antibodies (IRDye® 800CW Donkey anti-Rabbit IgG Secondary Antibody, LICORbio, 926-32213; or IRDye® 800CW Donkey anti-Mouse IgG Secondary Antibody, LICORbio, 926-32212) diluted 1:10,000 with TBS-T for 1 h at RT. Membranes were washed 3x15 min with TBS-T, rinsed with TBS (20 mM Tris pH 7.5, 150 mM NaCl) and imaged using LICORbio Odyssey M imager.

***C. elegans* dietary chimyl alcohol supplementation**

Animals were age-synchronized using hypochlorite treatment, and approximately 3,000 eggs were plated on either control plates (0.5% DMSO) or rescue plates (1 mM batyl or chimyl alcohol). The animals were cultured for 72 hours at 20°C. Upon collection, they were rinsed from the plates with M9 buffer and allowed to settle by gravity for five minutes to separate young progeny and eggs from Day 1 adults as effectively as possible. The M9 solution was aspirated down to the pellet and washed again with M9, followed by centrifugation at 1,000 x g for 30 seconds. Worm pellets were transferred to 1.5 mL Eppendorf tubes and flash-frozen in liquid nitrogen. To ensure proper solubilization of batyl or chimyl alcohol in RNAi, the batyl or chimyl alcohol was flash heated at 90°C for 15 seconds, then incubated with RNAi at 37°C for 15 minutes while shaking at 250 rpm.

**Fluorescent microscopy**

Confocal micrographs were acquired using a Leica SP8 confocal microscope (Leica) and Leica Application Suite X (LAS X) software (ver. 3.5.5). Live worms were mounted in M9 containing 200 mM levamisole and imaged at 40× and 63× magnifications with oil immersion. Imaging was performed in xyz acquisition mode, with each line imaged three times and averaged to reduce noise. Fluorophores were excited using 488 nm and 552 nm lasers with hybrid detectors, and laser power, range, and gain were adjusted based on the strain.

**Hydroxy acid LC-MS/MS analysis in *C. elegans***

For each sample, approximately 4000 *C. elegans* nematodes were thawed, and excess PBS was removed. A 100 µL mixture of 1:1 H_2_O:trifluoroethanol was added, followed by vortexing. Samples were then flash-frozen in quadruplicate, allowing thawing time between each freeze. The sample mixtures were transferred to ceramic bead tubes and homogenized using a TissueLyser II (4 cycles of 40 seconds at 30 Hz; cooled at 4°C for 5 minutes between cycles). Homogenized samples were transferred to new 1.5 mL tubes and mixed with 200 µL of 1:1 MeOH:EtOH containing 312 ng of d_4_-succinate and 10 µL of SPLASH II LipidoMIX^TM^ (Avanti, A83709), followed by vortexing. The samples were incubated on ice for 10 minutes. Next, 200 µL of water was added, mixed by pipetting, and incubated on ice for another 10 minutes. Samples were centrifuged at 16,000 ×g for 10 minutes at 4°C to pellet the solids.

The supernatants were transferred to Agilent Captiva EMR-Lipid 1 mL cartridges (Agilent 5190-1002). Cartridge elution was performed using a positive pressure manifold with ultra-pure nitrogen gas. The metabolite-containing flowthrough was collected in new 1.5 mL tubes. Cartridges were washed twice with 250 µL of 2:1:1 H_2_O:MeOH:EtOH, and the flowthroughs were collected in the same tubes. The lipid fractions were eluted by washing each cartridge twice with 600 µL of 2:1 MeOH:DCM, and the flowthroughs were collected in new 1.5 mL tubes. The collected fractions were evaporated until dry using a SpeedVac and stored at -80°C until analysis.

The metabolite-containing fraction was reconstituted in 100 µL of 70:20:10 ACN: H_2_O:MeOH. Samples were analyzed via LC-MS/MS on an Agilent 6595C triple quadrupole mass spectrometer (QqQ) using an InfinityLab Poroshell 120 HILIC-Z column (Agilent 683775-924, 2.7 µm, 2.1 × 150 mm). The column was maintained at 15 °C. The chromatography gradient began at 10% mobile phase A, comprised of 20 mM ammonium acetate in water (pH 9.3) and 5 µM of medronic acid, and 90% mobile phase B, comprised of 100% HPLC-grade ACN. The gradient proceeded as follows: starting at 10% mobile phase A, it increased to 22% over 8 min, then to 40% by 12 min, 90% by 15 min, and then held at 90% until 18 min before re-equilibration at 10% (held until 23 min). The flow rate was maintained at 0.4 mL/min for most of the run but increased to 0.5 mL/min from 19.1 min to 22.1 min to fully re-equilibrate the column between injections.

The UHPLC system was connected to an Agilent 6595C QqQ MS dual AJS ESI mass spectrometer. This method was operated in positive mode. The gas temperature was maintained at 200°C with flow at 14 L/min. The nebulizer was maintained at 50 PSI, sheath gas temperature at 375°C, and sheath gas flow at 12 L/min. VCap voltage was set to 3000V, iFunnel high-pressure RF was set to 150 V, and iFunnel low-pressure RF was set to 60 V. A dMRM inclusion list was used to individually optimize fragmentation parameters.

Data were collected in .d format and checked manually in Agilent MassHunter Qualitative Analysis and uploaded to MassIVE for data availability. For analysis, the data were then uploaded to Agilent MassHunter Quantitative Analysis for quantitation using relative internal standard calculations to calculate analyte concentrations. After manual inspection and integration, analyte concentration (ng/mL of reconstituted extract) was exported to .csv files.

**Untargeted lipidomics for *C. elegans***

For untargeted lipidomics, lipids were separated on an Agilent 1260 Infinity II UHPLC system coupled to an Agilent 6546 quadrupole time-of-flight Acquity, using a BEH C18 column (Waters 186009453, 1.7 µm 2.1 × 100 mm) at 50 °C with a VanGuard BEH C18 precolumn (Waters 18003975). The chromatographic gradient began at 85% mobile phase A, which consisted of 60:40 ACN:H_2_O with 10 mM ammonium formate and 0.1% formic acid, and 15% mobile phase B consisted of 9:1:90 ACN:H_2_O:IPA with 10 mM ammonium formate and 0.1% formic acid. The flow rate was 0.5 mL/min. The gradient increased to 30% mobile phase B during the next 2.4 min. The gradient then increased to 48% until 3 min, and then to 82% at 13.2 min. From 13.2 to 13.8 min, the gradient increased to 99%, and stayed at 99% until 16 min. At 16 min, re-equilibration to 15% mobile phase B began and was held until 20 min.

In negative mode, the gas temperature was maintained at 250 °C at a flow rate of 12 L/min. The sheath gas was maintained at 375 °C at a flow rate of 12 L/min. The nebulizer was set to 30 PSI. Vcap voltage was set to 4000 V, the skimmer was set to 75 V, the fragmentor was set to 190 V, and the octapole radiofrequency peak was set to 750 V. In positive mode, all mass spec settings were the same, except that the nebulizer was set to 35 PSI, and the sheath gas was maintained at 300 °C with a flow rate of 11 L/min. Reference masses used for positive mode were 121.05 and 922.01 *m/z*, and reference masses used for negative mode were 112.98 and 1033.99 *m/z*. For both ionization modes, the acquisition rate was 3 spectra/s, and the *m/z* range was 100–1700 *m/z*. For MS2 scans in both modes, the isolation width was set to narrow (1.3 *m/z*), the acquisition rate was 2 spectra/s, and the collision energy was fixed at 25 V. Precursors were excluded after 1 spectrum.

3 µL of each sample was injected in positive mode, and 5 µL was injected in negative mode. For data analysis, libraries of identified lipids were created using MS/MS spectra of 6 consecutive injections of pooled sample using iterative exclusion. Lipid annotation and library generation were performed using Agilent Lipid Annotator. Individual sample peak integration was completed using Agilent Profinder. Data normalization was performed in R using publicly available custom scripts (<https://github.com/RJain52/Multi-Tissue-Cold-Exposure-Lipidomics>).

**Mouse genotyping and sample collection**

Genotyping was confirmed by tail clipping through TransNetyx (Wild type Forward: GTTCTGGAAAATGATGTTGCCT; Reverse: AGTGCCATCCCATAATACCTCA; Reporter: CATACTCAGTGTCAACTCTAA; ACAD10 KO Forward: GGCTATTCTCAGGCAAAGGCT; Reverse: AGTGCCATCCCATAATACCTCA; Reporter: CTCAGGTCATCAGACTCTG)

Before sample collection, mice were fasted for 6 hours with access to water prior to euthanasia. Mice were anesthetized with isoflurane at 3.5% and euthanized by cervical dislocation. Blood was collected by cardiac puncture into sodium citrate buffer tubes (Covidien). Plasma was isolated by centrifugation (10 mins, 2000 x g, 4°C). Supernatants were transferred to new tubes and flash-frozen in liquid nitrogen. Plasma samples were stored at -80°C until the day of extraction. Mouse livers were harvested and flash-frozen in liquid nitrogen. Frozen livers were stored at -80°C until the day of extraction.

**LC-MS/MS Lipidomics**

*Lipidomics of* C.elegans*:* The lipid fraction was reconstituted in 50 µL of isopropanol. Lipids were separated on an Agilent 1260 Infinity II UHPLC system coupled to an Agilent 6546 quadrupole time-of-flight Acquity, using a BEH C18 column (Waters, 186009453, 1.7 µm 2.1 × 100 mm) at 50 °C with a VanGuard BEH C18 precolumn (Waters, 18003975). The chromatographic gradient began at 85% mobile phase A, which consisted of 60:40 ACN:H_2_O with 10 mM ammonium formate and 0.1% formic acid, and 15% mobile phase B consisted of 9:1:90 ACN:H_2_O:IPA with 10 mM ammonium formate and 0.1% formic acid. The flow rate was 0.5 mL/min. The gradient increased to 30% mobile phase B during the next 2.4 min. The gradient then increased to 48% until 3 min, and then to 82% at 13.2 min. From 13.2 to 13.8 min, the gradient increased to 99%, and stayed at 99% until 16 min. At 16 min, re-equilibration to 15% mobile phase B began and was held until 20 min.

In negative mode, the gas temperature was maintained at 250 °C at a flow rate of 12 L/min. The sheath gas was maintained at 375 °C at a flow rate of 12 L/min. The nebulizer was set to 30 PSI. Vcap voltage was set to 4000 V, the skimmer was set to 75 V, the fragmentor was set to 190 V, and the octapole radiofrequency peak was set to 750 V. In positive mode, all mass spec settings were the same, except that the nebulizer was set to 35 PSI, and the sheath gas was maintained at 300 °C with a flow rate of 11 L/min. Reference masses used for positive mode were 121.05 and 922.01 *m/z*, and reference masses used for negative mode were 112.98 and 1033.99 *m/z*. For both ionization modes, the acquisition rate was 3 spectra/s, and the *m/z* range was 100–1700 *m/z*. For MS2 scans in both modes, the isolation width was set to narrow (1.3 *m/z*), the acquisition rate was 2 spectra/s, and the collision energy was fixed at 25 V. Precursors were excluded after 1 spectrum.

3 µL of each sample was injected in positive mode, and 5 µL was injected in negative mode. For data analysis, libraries of identified lipids were created using MS/MS spectra of 6 consecutive injections of pooled sample using iterative exclusion. Lipid annotation and library generation were performed using Agilent Lipid Annotator. Individual sample peak integration was completed using Agilent Profinder. Data normalization was performed in R using publicly available custom scripts ([https://github.com/RJain52](https://github.com/RJain52/Multi-Tissue-Cold-Exposure-Lipidomics)).

*Lipidomics of mouse liver and plasma:* Lipids were extracted with a mix of isopropanol, water, and ethyl acetate (3:1:6) with 0.01% BHT and added internal standards: SPLASH Lipidomix II (10 µL/sample), oleoyl-L-carnitine d3 (2.33 pmol/sample), C18(plasm)-18:1(d9) (2.56 pmol/sample), and stearic acid d35 (90 nmol/sample). For external standard matrix PC C18(Plasm)-22:6 PC (2.56pmol/sample) and C16(Plasm)-18:1 PC (2.56pmol/sample) were used to assess retention time.

500 μL of extraction mix was added to 25 µL plasma or 19-22mg liver in ceramic bead tubes (Omni Int #19-627). Samples were shaken in a TissueLyzer (frequency 30/s, 30s) until fully homogenized, with a break (5min, 4°C) every 2 cycles. Lysates were incubated at -20°C for 10 min and then centrifuged (16,000 x g, 4°C, 10 min). 425 µL of supernatant was transferred to a microcentrifuge tube, the spin was repeated, and the subsequent 350 µL supernatant was transferred to a new tube. Extracts were evaporated by vacuum (40°C, 2 hr).

Lipidomics was performed by ultrahigh-performance liquid chromatography (Agilent 1290 Infinity II BioLC) on a ZORBAX C18 column (Agilent #959758-902) followed by triple quadrupole mass spectrometry (Agilent 6495C) using the chromatography gradient and dynamic multiple reaction monitoring (dMRM) settings outlined in^26^ (Agilent Application Note #5994-3747EN). Dried lipid extracts were resuspended in isopropanol (150 µL), transferred to injection vials with glass inserts (Agilent #5183-2086), and capped (Agilent #5185-5820). Liver samples were diluted 1:8 in isopropanol. Samples were maintained at 21°C in the multisampler, and 1 µL injections were run in a randomized order. Pooled matrix samples were used to update retention times from the published protocol. Peak integration was validated based on available data of isotope patterns; lysophospholipids with two resolved peaks were annotated “a” or “b.” Peaks were smoothed via the Quadratic/Cubic Savitzky-Golay method and then normalized to the relative peak abundance of a representative internal standard using MassHunter Quantitative Analysis (Agilent). Peaks that had an equal or higher abundance in a no-sample blank were removed. Raw data files were converted to mzML and uploaded to MassIVE (#MSV000096321).

*Targeted oxylipin analysis of mouse plasma*: The extraction mix was composed of MeOH containing 0.02% (w/v) BHT and 1.5 ng per sample of each of the following internal standards: 14(15)-EET-d11, TXB2-d9, LTB3-d4, 9-HODE-d4, PGD2-d4, PGE2-d4, 20-HETE-d6, LXA4-d5, 14(15)-DHET-d11, AA-d11, EPA-d5, DHA-d5, 12(13)-DiHOME-d4, and Maresin-1-d5. 1 mL of the extraction mix was added to 50 µL of plasma, followed by vortexing. Samples were incubated at -80°C for one hour. After incubation, samples were centrifuged at 16,000 ×g for 10 minutes at 4°C to pellet the solids. Supernatant was transferred to new 1.5 mL tubes. Samples were evaporated until dry using a SpeedVac, then stored at -80°C until analysis, at which point they were resuspended in 50 µL of 1:1 MeOH:H_2_O.

Oxylipin extracts were analyzed using an Agilent Infinity II LC coupled to an Agilent 6495c triple quadrupole mass spectrometer. The LC was equipped with an Agilent RRHD Eclipse Plus C18 column (2.1 × 150 mm, 1.8 µm) with an Agilent Eclipse Plus guard column (2.1 × 5 mm, 1.8 µm) kept at 50°C. Mobile phase A consisted of 0.1% acetic acid and mobile phase B was ACN:IPA (90:10, v/v). The LC gradient was as follows: start at 15% B to 33% at 3.5 min, to 38% B at 5.5 min, to 42% B at 7 min, to 48% B at 9 min, to 65% B at 15 min, to 75% B at 17 min, to 85% B at 18.5 min, to 95% B at 19.5 min, and finally to 15% B at 21 min, held until 26 min at a constant flow of 0.350 mL/min. Multi-sampler was kept at 4°C and blank injections were run between every sample. Samples were injected in random order and within 48 hrs of extraction.

MS analysis was in negative ionization with the following parameters: drying gas at 290°C at 10 L/min, nebulizer at 35 psi, sheath gas at 350°C at 11 L/min, capillary voltage at 3500 V and nozzle voltage at 1000 V. A dynamic multi-reaction monitoring (dMRM) method was used for analysis. Data were collected and analyzed using the Agilent MassHunter Suite (v10.1). Automated peak picking was used to integrate peaks with a signal-to-noise ratio > 3.0 and retention times within 0.30 min of the expected retention time. All peaks were then manually assessed for appropriate shape. Any samples with peak height less than background were removed. Compounds present in at least 60% of samples were reported in the final dataset in units of “ng lipid per mL plasma” after normalization to the appropriate internal standard.

**Hepa 1-6 cell sample collection**

For the batyl alcohol Hepa1-6 treatments, cells were grown to confluence, then treated with 20 μM batyl alcohol or an equivalent volume of ethanol as a vehicle control for 16 h. At 16 h, media was collected and flash frozen. Then cells were washed 3x with sterile PBS, scraped to disrupt adhesion, and transferred to an Eppendorf tube for flash freezing until day of the extraction.

**Targeted LC-MS/MS measurement of platelet-activating factor**

For platelet-activating factor measurements in cell media and human plasma, an extraction solvent was prepared by adding 70 µL of D4-PAF-C16 to 27 mL of methanol. 1 mL of extraction solvent was added to 200 μL of cell media or 100μL of human plasma. Samples were vortexed until fully homogenized, then allowed to incubate at -80°C for one hour. After incubation, samples were centrifuged at 16000 xg for 10 minutes at 4°C. The supernatant was transferred to a separate microcentrifuge tube and evaporated to dryness. Samples were reconstituted in 100 µL of 70:20:10 ACN:H_2_O:MeOH. PAF-16 was measured with the HILIC-Z gradient outlined above.

Data from human studies is held in the Data for Indigenous Implementations, Interventions, and Innovations Tribal Data Repository. Access requires review and approval by the D4I TDR Data Access and Use Committee (DAUC) and, where applicable, by the originating Tribal Nation, in accordance with the governance templates and data-use agreements maintained at [https://d4itdr.org/resources/](https://urldefense.com/v3/__https:/d4itdr.org/resources/__;!!MznTZTSvDXGV0Co!GKwMWGUXQ28wCa8hS9UO5zKBvWiqJpGGmapP9DMN4q-x-xvlmakllHbIuoXEj9hoJ2Hg7d2PNKl6Em_s96ooY9Jm7OxHnA$). These materials are licensed under CC BY-NC 4.0 and outline the terms under which participating Tribes and Indigenous communities retain authority over how their data are accessed, used, and shared."

**LC-MS/MS proteomics**

Hepa 1-6 cells were cultured under standard conditions according to ATCC guidelines with DMEM with 10% FBS, with the addition of primocin as an antimicrobial. Cells were harvested at 100% confluence by scraping following two washes with ice-cold PBS. Cell pellets were lysed in 200 µL of urea lysis buffer (8 M urea, 100 mM ammonium bicarbonate, 1x protease inhibitor cocktail, and 50 mM TEAB, pH ~8) and disrupted by bead homogenization in ceramic bead tubes with 4 cycles of 20 sec disruption at 6.0m/s. Lysates were clarified by centrifugation 14,000 × g, 10 min, 4 °C, and total protein concentration in the resulting supernatant was determined by Pierce BCA assay using a BSA standard curve.

For each sample, 100 µg of total protein was reduced with 5 mM DTT, 56 °C, 30 min and alkylated with 15 mM iodoacetamide, room temperature, 30 min, dark. Samples were diluted with 100 mM ammonium bicarbonate to reduce the urea concentration below 2 M and digested with sequencing-grade modified trypsin (Promega) at a 1:10 (enzyme:protein) ratio for 16 hr at 37 °C with gentle agitation. Digestion was quenched by acidification to pH>3 with 10% formic acid. Peptides were desalted using Pierce C18 spin columns per the manufacturer's protocol, eluted in 70% acetonitrile/0.1% TFA, dried by SpeedVac, and reconstituted in 20μL of 2% acetonitrile, 0.1% formic acid prior to LC-MS/MS analysis.

Digested peptides were separated by reverse-phase nanoflow liquid chromatography using an Agilent 1100 Nanopump fitted with an Easy-Spray column (15 cm x 75μm, packed with Pepmap RSLC C18, 3μm, ThermoFisher). Peptides were eluted over a 230-min linear gradient from 5% to 30% mobile phase B (mobile phase A: 0.1% formic acid in water; mobile phase B: 0.1% formic acid in acetonitrile) at a flow rate of 300 nL/min. Eluted peptides were analyzed by a Thermo Scientific Orbitrap Fusion Lumos Tribrid mass spectrometer operated in parallel reaction monitoring (PRM) mode. A scheduled PRM method was built in Skyline targeting proteotypic peptides unique to human ACAD10, selected on the basis of in silico digestion and uniqueness filtering against the UniProt proteome. Precursor ions were isolated in the quadrupole 0.7 Da isolation window and fragmented by higher-energy collisional dissociation (HCD, normalized collision energy 38%), with product ions detected in the Orbitrap at a resolution of 30,000. A retention-time scheduling window of 5 min was applied around predicted elution times determined from a preceding discovery-mode LC-MS/MS run of a pooled sample digest.

Raw PRM data files were imported into Skyline (version 25.1, MacCoss Lab, University of Washington) for peak detection, manual review of chromatographic peak boundaries, and extraction of integrated peak areas for the top 6 most intense, interference-free product ions per targeted peptide. Peak areas were normalized to total peptide amount injected/total ion current to generate relative quantitative values across samples. Peptide-level values for each protein target were used to generate protein-level abundance estimates for ACAD10 in each sample. Peptide identifications underlying the PRM assay were confirmed by comparison of fragment ion retention times and relative intensity ratios to a reference spectral library generated from a discovery-mode DDA run of the pooled digest, searched against the UniProt reference proteome using Proteome Discoverer version 3.2 with 10 ppm precursor mass tolerance filtered to 1% false discovery rate.

3. **Quantification and Statistical analysis**

**Lipidomic Data analysis**

To compare global lipid composition in *C. elegans* or mouse liver samples by class, lipids were grouped:

| Lipid class | Lipids |
| --- | --- |
| Phosphoglycerolipids | Lysophosphatidylcholine (LPC), lysophosphoatidylethanolamine (LPE),  Lysophosphatidylinositol (LPI),  phosphatidic acid (PA), phosphatidylcholine (PC), phosphatidylethanolamine (PE), phosphatidylserine (PS), phosphatidylglycerol (PG), phosphatidylinositol (PI), phosphatidylinositol phosphate (PIP1), Ether lipid phosphatidylethanolamine (EtherPE*), cardiolipin (CL), phosphatidylethanols (PEtOH). |
| Sphingolipids | Sphingomyelin (SM), ceramides (Cer), dihydroceramide (dhCer), glycosphingolipid 3 (GM3), hexosylceramide (HexCer), Sulfoglycosphingolipids (SHexCer) |
| Cholesteryl esters | Cholesteryl ester (CE), methyl-cholesteryl ester (methyl-CE), dimethyl-cholesteryl ester (dimethyl-CE) |
| Glycerolipids | Diacylglycerol (DAG), triacylglycerol (TAG) |
| Fatty acids and metabolites | Acylcarnitine (AC), fatty acids (FA), Fatty Acyl Esters of Hydroxy Fatty Acid (FAHFA) |

* Under global targeted lipidomics experimental set up ether lipids cannot be confidently assigned to ether or ether-vinyl linkage lipids.

To compare phosphoglycerolipid acyl chain lengths and saturation, phospholipids with identified acyl chain lengths were included into analysis (excluding lipids containing two acyl chains without neutral loss data to identify the individual acyl chains). Ether lipid chains were excluded from the analysis.

**Heatmap generation for *C. elegans* and mouse liver ether lipids**

To present the data as fold change from the WT control group, adjustment of data had to be made for lipids below the lower limit of detection resulting in 0 values being assigned. 95% of the lowest detected value from all lipids was added to all the data values before the data was adjusted as fold change from the WT control group average. This conversion was only applied to the heatmap data. Data was presented as a heatmap using GraphPad Prism 10, showing individual replicate values within each group.

**Bioinformatics**

Homologs of ACAD10 kinase, phosphatase and dehydrogenase domains were fetched by querying the Uniprot database with Pfam domains PF01636, PF00702, PF02770, respectively, using only representative proteomes estimated to be at least 85% complete as per BUSCO score ^65^. Only those homologs were retained for which ACAD10 or ACAD11 were the top BLAST hit in a reverse search versus the human Uniprot database. AGPS homologs were fetched likewise, querying with the Pfam domain PF02913. The taxonomy dendrogram annotated with ACAD and AGPS occurrence histograms was visualized in iToL ^66^. Sequence alignments were created with ClustalW ^67^ and visualised in ESPript ^68^.

**Statistical analysis**

For analysis of lipid level change depending on diet and genotype, two-way ANOVA with Šídák multiple comparisons test was performed for each lipid species using Prism 10 comparing WT control vs ether lipid supplemented and Acad10 KO (or *acds-10 -/-*) control vs ether lipid supplemented. For analysis of lipid level change depending on genotype, unpaired two-tailed t-tests were performed using Prism 10. All results are presented as mean +/- SD with individual values shown.

For the mouse plasma oxylipin data volcano plot, changes in plasma lipid levels were analyzed using two-tailed Students t-tests. The resulting p-values were transformed by -log_10_ and plotted against magnitude of change between Acad10 KO and WT mice transformed by log_2_. Values with statistical significance (p<0.05) were highlighted.

For *C. elegans* lifespan analysis Prism 10 was used, p-values were calculated using the long-rank (Mantel-Cox) method.

During the preparation of this work, the authors used ChatGPT to check grammar, spelling and improve sentence structure. After using this tool/service, the author(s) reviewed and edited the content as needed and take full responsibility for the content of the publication.

4. **Key resources table**

| REAGENT or RESOURCE | SOURCE | IDENTIFIER |
| --- | --- | --- |
| Antibodies | | |
| *C. elegans* tubulin | Sigma-Aldrich | T6074 |
| ACDS-10 | This study | N/A |
| Bacterial and virus strains | | |
| BL21 (DE3) *E. coli* | Orth Lab | N/A |
| Rosetta (DE3) *E. coli* | Orth Lab | N/A |
| OP50 *E. coli* | Orth Lab | N/A |
| HT115 *E. coli* | Orth Lab | N/A |
| Biological samples |  |  |
| Human plasma | Tribal Data Repository (D4I TDR) | N/A |
| Chemicals, peptides, and recombinant proteins | | |
| ATP | Sigma-Aldrich | A2383 |
| [γ-^32^P]ATP | Revvity | BLU002Z001MC |
| AMP-PNP | Sigma-Aldrich | 10102547001 |
| DTT | Goldbio | DTT25 |
| Water | Honeywell | LC365 |
| Methanol | Honeywell | LC230 |
| Ethanol | Sigma-Aldrich | 91683 |
| Trifluoroethanol | Sigma-Aldrich | 91683 |
| Dichloromethane | Sigma-Aldrich | 650463 |
| Acetonitrile | Honeywell | LC015 |
| Ammonium formate | Honeywell | 55674 |
| Ethyl acetate | Sigma-Aldrich | 650528 |
| Butylated hydroxytoluene | Sigma-Aldrich | B1378 |
| Isopropanol | Fisher Scientific | A461 |
| Formic acid | Fisher Scientific | A11710X1 |
| InfinityLab deactivator additive | Agilent | 5191-4506 |
| Chimyl alcohol/1-O-hexadecyl-*rac*-glycerol | Santa Cruz | 6145-69-3 |
| 1-O-hexadecyl-2-palmitoyl-*rac*-glycerol | Biosynth | FH172161 |
| batyl alcohol/1-O-octadecyl-*rac*-glycerol | Sigma Aldrich | B402-1G |
| 1-O-hexadecyl-2-acetyl-*sn*-glycerol | Enzo Life Sciences | #50-201-0570 |
| DL-α -palmitin/1-Monohexadecanoyl-*rac*-glycerol | Sigma-Aldrich | M1640 |
| 1-Oleoyl-2-acetyl-*sn*-glycerol | Sigma Aldrich | 495414 |
| 1-Palmitoyl-2-Lauroyl-*rac*-glycerol | Cayman Chemical | 26875 |
| d_4_-PAF-C16 | Cayman Chemical | 360900 |
| Coenzyme A sodium salt | Sigma-Aldrich | C3144 |
| acetyl coenzyme A sodium salt | Sigma-Aldrich | A2056 |
| n-propionyl coenzyme A lithium salt | Sigma-Aldrich | P5397 |
| malonyl coenzyme A lithium salt | Sigma-Aldrich | M4263 |
| succinyl coenzyme A sodium salt | Sigma-Aldrich | S1129 |
| butyryl coenzyme A lithium salt | Sigma-Aldrich | B1508 |
| crotonyl coenzyme A | Sigma-Aldrich | 28007 |
| DL-3-Hydroxy-3-methylglutaryl (HMG) coenzyme A | Sigma-Aldrich | H6132 |
| hexanoyl coenzyme A trilithium salt | Sigma-Aldrich | H2012 |
| 3’-dephospho coenzyme A | Cayman Chemical | 27390-1 |
| SPLASH II LipidoMIX^TM^ Mass spec standard | Avanti | A83709 |
| oleoyl-L-carnitine d_3_ | Cayman Chemical | 26578 |
| d_4_-succinate | Sigma Aldrich | 293075 |
| C18(Plasm)-22:6 PC | Avanti | 852472 |
| C16(Plasm)-18:1 PC | Avanti | 852478 |
| stearic acid d35 | Cayman Chemical | 9003318 |
| 14(15)-EET-d11 | Cayman Chemical | 10006410 |
| TXB2-d9 | Cayman Chemical | 9002563 |
| LTB4-d4 | Cayman Chemical | 320110 |
| 9-HODE-d4 | Cayman Chemical | 338410 |
| PGD2-d4 | Cayman Chemical | 312010 |
| PGE2-d4 | Cayman Chemical | 314010 |
| 20-HETE-d6 | Cayman Chemical | 390030 |
| LXA4-d5 | Cayman Chemical | 10007737 |
| 14(15)-DHET-d11 | Cayman Chemical | 10008040 |
| AA-d11 | Cayman Chemical | 10006758 |
| EPA-d5 | Cayman Chemical | 10005056 |
| DHA-d5 | Cayman Chemical | 10005057 |
| 12(13)-DiHOME-d4 | Cayman Chemical | 10009994 |
| Maresin-1-d5 | Cayman Chemical | 21823 |
| Deposited data | | |
| Raw TLC files and lipidomics/proteomics exports | Unpublished Mendeley Data | <https://data.mendeley.com/preview/csctm9nz8v?a=f0831168-4d78-4403-bc50-f4f0c7e57c60> |
| Raw LC/MS data | MassIVE | [https://massive.ucsd.edu/ProteoSAFe/static/massive.jsp?redirect=auth](https://urldefense.com/v3/__https:/massive.ucsd.edu/ProteoSAFe/static/massive.jsp?redirect=auth__;!!MznTZTSvDXGV0Co!GKwMWGUXQ28wCa8hS9UO5zKBvWiqJpGGmapP9DMN4q-x-xvlmakllHbIuoXEj9hoJ2Hg7d2PNKl6Em_s96ooY9LyDC-RPg$)  (MSV000103144; MSV000103078; MSV000097576; MSV000097543; MSV000103165) |
| Experimental models: Cell lines | | |
| Hepa 1-6 parental (WT) | ATCC | CRL-1830 |
| Hepa 1-6 Acad10-KO | This study | N/A |
| Hepa 1-6 Acad11-KO | This study | N/A |
| Hepa 1-6 Acad10/Acad11-DKO | This study | N/A |
| Hepa 1-6 WT pQCXIP EV | This study | N/A |
| Hepa 1-6 WT pQCXIP ACAD10-Flag WT | This study | N/A |
| Hepa 1-6 Acad10-KO pQCXIP EV | This study | N/A |
| Hepa 1-6 Acad10-KO pQCXIP ACAD10-Flag WT | This study | N/A |
| Hepa 1-6 Acad10-KO pQCXIP ACAD10-Flag D48A | This study | N/A |
| Hepa 1-6 Acad10-KO pQCXIP ACAD10-Flag D483A | This study | N/A |
| Hepa 1-6 Acad10-KO pQCXIP ACAD10-Flag D1040A | This study | N/A |
| Hepa 1-6 Acad10-KO pQCXIP 41-1059 ACAD10-Flag (ΔMTS) | This study | N/A |
| Hepa 1-6 Acad10-KO pQCXIP Flag-41-1059 ACAD10 + 721-780 ACAD11 (PTS) | This study | N/A |
| Hepa 1-6 Acad10-KO pQCXIP 1-270 ACAD10 + 26-357 ACAD11 + 609-1059 ACAD10-Flag (A11KD) | This study | N/A |
| Experimental models: Organisms/strains | | |
| *Acad10*^em1(IMPC)Tcp^ C57BL/B6J mice | MMRRC | 066557-UCD |
| *C. elegans* N2 (WT) | CGC | N2 |
| *C. elegans* Δacds-10 | SUNY biotech | VTA01 (syb4573) |
| *C. elegans* mcherry::acds-10 | SUNY biotech | PHX9793 |
| *C. elegans* *dhs-3p::DHS-3::GFP* | CGC | LIU1 |
| *C. elegans* mcherry::acds-10 x *dhs-3p::DHS-3::GFP* | This study | N/A |
| *C. elegans* *eft-3p::3XFLAG::GFP::SKL::unc-54 3'UTR* | CGC | WBM1177 |
| *C. elegans* mcherry::acds-10 x *eft-3p::3XFLAG::GFP::SKL::unc-54 3'UTR* | This study | N/A |
| *C. elegans* acds-10 D9A | SUNY biotech | PMD26 (*syb8910*) |
| *C. elegans* acds-10 D424A | SUNY biotech | PMD27 (*syb8950*) |
| *C. elegans* acds-10 D966A | SUNY biotech | PMD28 (*syb9051*) |
| Oligonucleotides | | |
| GTTCTGGAAAATGATGTTGCCT | Mouse wild type genotyping: forward | N/A |
| AGTGCCATCCCATAATACCTCA | Mouse wild type genotyping: reverse | N/A |
| CATACTCAGTGTCAACTCTAA | Mouse wild type genotyping: reporter | N/A |
| GGCTATTCTCAGGCAAAGGCT | mAcad10-KO genotyping: forward | N/A |
| AGTGCCATCCCATAATACCTCA | mAcad10-KO genotyping: reverse | N/A |
| CTCAGGTCATCAGACTCTG | mAcad10-KO genotyping: reporter | N/A |
| Recombinant DNA | | |
| pQCXIP EV | From Jack Dixon lab | N/A |
| pQCXIP ACAD10-Flag WT | This study | N/A |
| pQCXIP ACAD10-Flag D48A | This study | N/A |
| pQCXIP ACAD10-Flag D483A | This study | N/A |
| pQCXIP ACAD10-Flag D1040A | This study | N/A |
| pQCXIP 41-1059 ACAD10-Flag (ΔMTS) | This study | N/A |
| pQCXIP Flag-41-1059 ACAD10 + 721-780 ACAD11 (PTS) | This study | N/A |
| pQCXIP 1-270 ACAD10 + 26-357 ACAD11 + 609-1059 ACAD10-Flag (A11KD) | This study | N/A |
| ppSUMO EV (modified pET-28a containing an N-terminal 6X-His tag followed by the yeast Sumo (smt3) CDS) | From Jack Dixon lab | N/A |
| ppSUMO ACDS-10 209-568 WT | This study | N/A |
| ppSUMO ACDS-10 209-568 D424A | This study | N/A |
| ppSUMO ACDS-10 1-212 WT | This study | N/A |
| ppSUMO ACDS-10 1-212 D9A | This study | N/A |
| ppSUMO *S. raichei* (A0A060NI31_9BURK) ACAD10 kinase | This study | N/A |
| ppSUMO *Aggregiococcus sp*. (A0A6I2GPB5_9BACT) ACAD10 kinase | This study | N/A |
| ppSUMO M. pacifica (A0A4S2HBA5-9PROT) ACAD10 kinase | This study | N/A |
| ppSUMO C. aerophila (I0I2H8_CALAS) ACAD10 kinase | This study | N/A |
| ppSUMO Aeromicrobium sp (A0A4V2NSZ3_9ACTN) ACAD10 kinase | This study | N/A |
| ppSUMO M. xanthus (Q1CZY9_MYXXD) ACAD10 kinase | This study | N/A |
| ppSUMO C. thermarum (WP_007504381.1) ACAD10 kinase WT | This study | N/A |
| ppSUMO C. thermarum ACAD10 kinase R61A | This study | N/A |
| ppSUMO C. thermarum ACAD10 kinase E78A | This study | N/A |
| ppSUMO C. thermarum ACAD10 kinase D214A | This study | N/A |
| ppSUMO C. thermarum ACAD10 kinase K216A | This study | N/A |
| ppSUMO C. thermarum ACAD10 kinase N219A | This study | N/A |
| ppSUMO C. thermarum ACAD10 kinase D234A | This study | N/A |
| ppSUMO C. thermarum ACAD10 kinase E236A | This study | N/A |
| ppSUMO human ACAD10 41-617 WT | This study | N/A |
| ppSUMO human ACAD10 41-617 D48A | This study | N/A |
| ppSUMO human ACAD10 41-617 D483A | This study | N/A |
| ppSUMO human ACAD10 41-617 D48A+D483A | This study | N/A |
| ppSUMO ACAD11 11-356 WT | This study | N/A |
| ppSUMO ACAD11 11-356 D242A | This study | N/A |
| ppSUMO *E. Coli* DgkA | This study | N/A |
| Software and algorithms | | |
| ImageJ | (Schneider et al., 2012)^69^ | <https://imagej.nih.gov/ij/> |
| PyMol | Schrödinger, Inc. | N/A |
| DaliLite server | (Holm and Rosenstrom, 2010)^70^ | <http://ekhidna2.biocenter.helsinki.fi/dali/> |
| UniProt | UniProt Consortium ^71^ | <https://www.uniprot.org> |
| iToL | (Letunic and Bork, 2024) ^66^ | <https://itol.embl.de> |
| ClustalW | (Sievers and Higgins, 2021) ^72^ | <https://www.ebi.ac.uk/jdispatcher/msa/clustalo> |
| ESPript | (Gouet, Robert and Courcelle, 2003)^73^ | <https://espript.ibcp.fr/ESPript/ESPript> |
| BLAST server | (Altschul et al., 1990)^74^ | <https://blast.ncbi.nlm.nih.gov/Blast.cgi> |
| MAFFT server | (Katoh and Standley, 2013)^75^ | <https://mafft.cbrc.jp/alignment/software/> |
| Pfam database | (Finn et al., 2016)^76^ | <http://pfam.xfam.org/> |
| GraphPad Prism 10 | N/A | <https://www.graphpad.com/scientific-software/prism/> |
| Adobe Illustrator 2026 | N/A | <https://www.adobe.com/products/illustrator.html> |
| Phenix | (Adams et al., 2010)^62^. | <https://www.phenix-online.org/> |
| Coot | (Emsley et al., 2010)^61^ | <https://www2.mrc-lmb.cam.ac.uk/Personal/pemsley/coot> |
| CCP4 7.1 | (Winn et al., 2011)^77^ | <https://www.graphpad.com/scientific-software/prism/> |
| AlphaFold server | (Abramson et al., 2024)^78^ | <https://alphafoldserver.com> |
| Other | | |
| Silica TLC plates (Silica gel 60 F₂₅₄, L × W 20 cm × 20 cm, glass support) | Supelco | 1.05715 |
| Captiva EMR-Lipid 1 mL cartridges | Agilent | 5190-1002 |
| Standard chow diet | Formulab Diet | 5008 |
| High-energy diet | Formulab Diet | 5015 |
| Standard chow diet | Envigo | TD.00606 |
